# Internal Ribosome Entry Sites act as Effector Domain in linear and circular antisense long non-coding SINEUP RNAs

**DOI:** 10.1101/2023.05.25.542260

**Authors:** Sabrina D’Agostino, Abraham Tettey-Matey, Massimiliano Volpe, Bianca Pierattini, Federico Ansaloni, Pierre Lau, Carlotta Bon, Omar Peruzzo, Clarissa Braccia, Andrea Armirotti, Margherita Scarpato, Devid Damiani, Valerio Di Carlo, Laura Broglia, Elias Bechara, Gian Gaetano Tartaglia, Piero Carninci, Claudio Santoro, Francesca Persichetti, Luca Pandolfini, Stefano Espinoza, Silvia Zucchelli, Remo Sanges, Stefano Gustincich

**Author notes:** corresponding author, Stefano Gustincich, Deputy Director for Life Sciences Director – Central RNA Laboratory Istituto Italiano di Tecnologia (IIT) via Melen 83, Genoa (GE) 16152 - Italy. equally contributed to the paper, alphabetical order (Sabrina D’Agostino, Abraham Tettey-Matey). This work is dedicated to the memory of Silvia Zucchelli.

## Abstract

SINEUPs are antisense long non-coding RNAs that enhance translation of overlapping sense mRNAs through the activity of two domains: a <u>SINE</u>B2 sequence <u>UP-</u>regulating translation (Effector Domain, ED) and an antisense region providing target specificity (Binding Domain, BD). In this study, we demonstrate that the invSINEB2 sequence from the natural SINEUP *AS Uchl1* RNA is an Internal Ribosomal Entry Site (IRES) when acting in *cis* and that known viral and cellular IRES sequences can act as Effector Domain in synthetic SINEUPs.

To identify natural IRES-containing, non-coding RNAs with SINEUP-like activity, we focused on circular RNAs showing that the non-coding *circ5533*, transcribed from the *c-myc locus*, enhances endogenous protein expression of its target *PX Domain Containing Serine/Threonine Kinase Like* (*Pxk)* by increasing mRNA association to polysomes.

In summary, this study shows that natural and synthetic SINEUPs include linear and circular transcripts with an embedded IRES sequence as ED.

## Introduction

The ability of organisms, tissues and cells to maintain a condition of equilibrium in challenging, stressful environments, requires a coordinated effort in modulating gene expression. By tweaking key molecular steps, homeostatic responses mainly entail moderate changes in RNA and protein levels that have large consequences on the well-being of living entities.

The non-coding portion of the genome has provided a plethora of natural regulatory RNAs to meet these challenges. On the one hand, microRNAs (miRNAs) reduce the level of protein expression of many genes by inhibiting translation and/or accelerating mRNA degradation. However, miRNAs reduce modestly the mean expression of most targeted proteins, leading to the current model about their function in decreasing protein expression noise in physiological and stress conditions [1]. On the other hand, SINEUPs (<u>SINE</u>B2 sequence to <u>UP</u>-regulate translation), a class of antisense long-non-coding RNAs (lncRNAs), increase the expression of their sense protein-encoding genes by enhancing translation. The representative member of natural SINEUPs is *AS Uchl1*, a lncRNA antisense to the mouse orthologue of the human ubiquitin C-terminal hydrolase L1 (*Uchl1*) gene. Upon stress, *AS Uchl1* promotes the association of *Uchl1* mRNA to polysomes and the consequent higher UCHL1 protein expression within the physiological range in conditions of attenuation of the cap-dependent translation [2]. SINEUP activity depends on the combination of two RNA domains: an embedded Transposable Element (TE), the mouse SINEB2 sequence in inverted orientation (invSINEB2), which acts as Effector Domain (ED), determining the up-regulation of target mRNA translation [2,3] and an overlapping antisense region, defined as Binding Domain (BD), that drives SINEUP specificity.

Additional mouse and human natural SINEUPs lncRNAs are transcribed from genomic *loci* with partially overlapping sense/antisense transcript pairs organized in a head-to-head conformation (Figure 1a) [2,3], suggesting that SINEUPs can represent a general class of regulatory RNAs.

**Figure 1.**
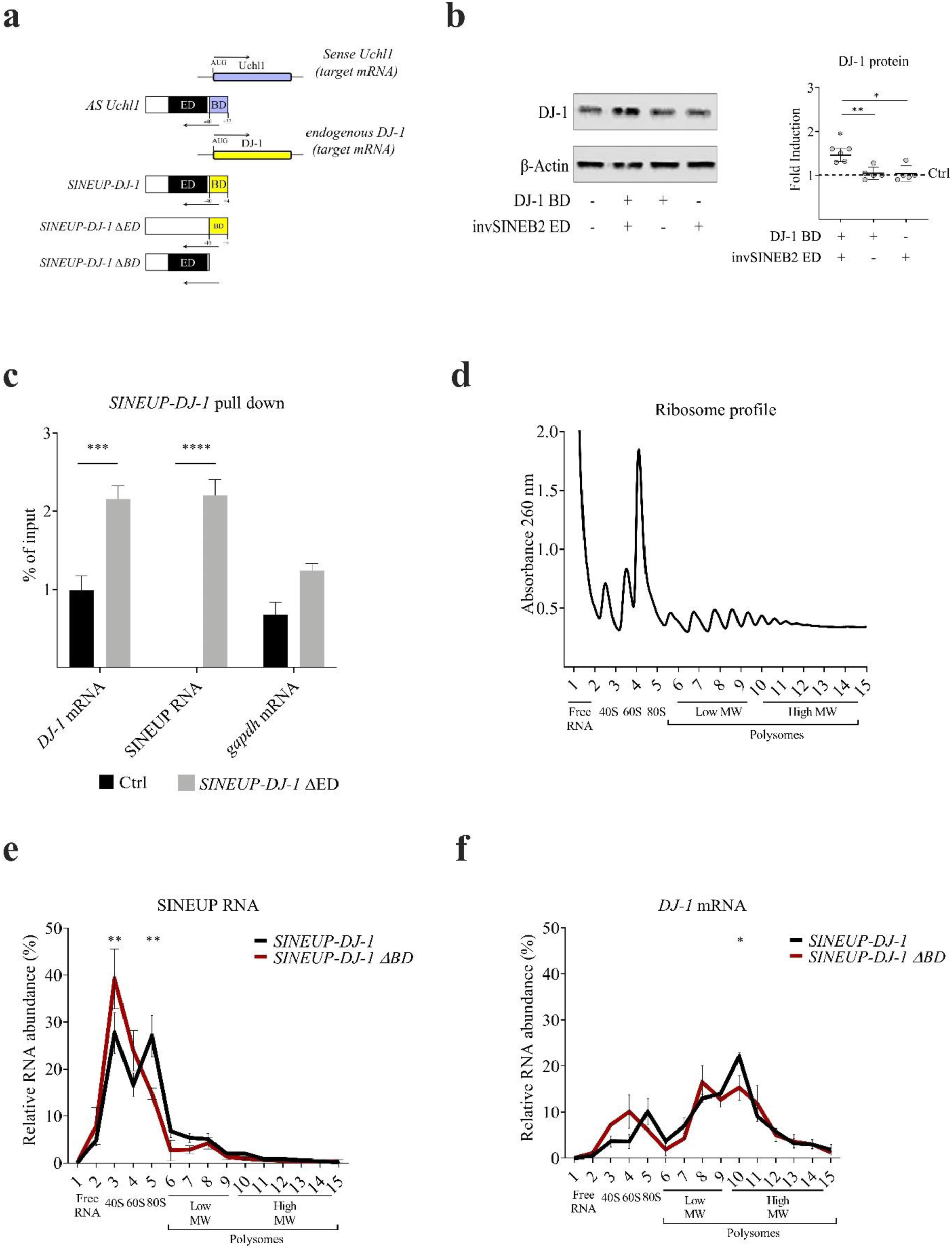
*SINEUP-DJ-1* increases DJ-1 protein translation. **a)** *AS Uchl1* and *SINEUP-DJ-1* targeting sense mRNAs. Natural antisense *Uchl1* comprises a BD (violet) pairing at the position −40/+32 (with A of the start codon placed at position +1) of sense *Uchl1* (target mRNA), a SINEB2 element acting as ED (black), and additional nts (white) from the *AS Uchl1* RNA backbone. Synthetic *SINEUP-DJ-1* partially overlaps *DJ-1* mRNA (yellow) in an antisense orientation; its activity relies on the presence of a BD (yellow) targeting the mRNA of interest (position −40/+4) and a SINEB2 element, in an inverted orientation (invSINEB2), acting as ED (black). *SINEUP-DJ-1 ΔED* lacks the ED (invSINEB2). *SINEUP-DJ-1 ΔBD* is a canonical synthetic SINEUP without an overlapping BD. **b)** Representative western blot image with reassuming graph. *SINEUP-DJ-1* constructs were transfected in HEK293T cells and tested as described in the Methods. Compared to the empty control, canonical *SINEUP-DJ-1* (with both DJ-1 BD and invSINEB2 ED) significantly increased endogenous DJ-1 protein levels. In contrast, SINEUP RNAs lacking BD or ED did not show any activity. Protein fold induction was quantified through western blot using anti-DJ-1 Ab. β-actin was used to normalize. Fold inductions were normalized to empty pCS2^+^ vector (Ctrl=1; dotted line). Plot reports mean ± SD of 5 replicates. *p <0.05 and **p <0.01 analyzed with ordinary one-way ANOVA (Holm-Sidak’s multiple comparisons test) were considered significant. **c)** *SINEUP-DJ-1* RNA Pull Down (RPD). *SINEUP-DJ-1* was purified using a 115 nts biotinylated RNA probe. As a negative control, RPD was performed in the absence of probes. qRT-PCR results show the enrichment of *SINEUP-DJ-1* and *DJ-1* mRNA upon *SINEUP-DJ-1* RPD. Data are shown as means % enrichment over input relative to the negative control ± SD of 5 replicates. Significant *p <0.05 and **p <0.01 values were analyzed with unpaired t-test. **d)** Polysome profile. Total lysates from control and treated cells were fractionated on sucrose gradients. Gradient fractions were separated through a UV detector continuously measuring the solution absorbance (260 nm) and plotting a chart showing the polysome profile of the gradient. **e)** Relative RNA abundance. The relative distribution (in %) of *SINEUP-DJ-1* and *SINEUP ΔBD* RNAs was analyzed by RT-qPCR in each of the 14 gradient fractions. RNA distributions were reported as the mean ± SD of 3 replicates. **f)** Relative *DJ-1* mRNA abundance. The relative distribution (in %) of *DJ-1* mRNA was analyzed, in the presence of *SINEUP-DJ-1* or *SINEUP ΔBD*, by RT-qPCR in each of the 14 gradient fractions. RNA distributions were reported as the mean ± SD of 3 replicates.

Artificial SINEUPs can be synthesized by designing antisense BD sequences for a gene of interest to specifically drive the enhancing activity of the invSINEB2 element to ectopically expressed mRNAs as well as endogenous ones [3–5]. The target site (TS) is usually located at the 5’ untranslated region (5’UTR) of the mRNA and can include the AUG translation initiation site. The target gene’s protein expression increases by 1.5 to 2.5 folds, making SINEUPs an ideal tool to perturb gene expression *in vivo* within the physiological range and to treat unmet human diseases caused by haploinsufficiency [4–6].

The molecular mechanism of natural and synthetic SINEUP RNAs activity is still unclear. While SINE sequences, and TEs at large, were previously considered to be “junk”, they are now known to play pivotal roles in shaping genome diversity and transcriptome complexity [7,8]. TEs comprise a significant fraction of lncRNA sequences [9,10] that may exert their function acting as independent domains [2,3,11,12] through specific RNA-RNA and RNA-protein interactions [13,14]. A combination of chemical footprinting, NMR analysis, and functional studies showed that the apical Stem Loop 1 (SL1) of the invSINEB2 sequence from *AS Uchl1* is required for activity [15]. Nevertheless, how a SINE sequence does regulate translation remains unknown.

The translation machinery is extremely energy-consuming [16] for a cell, which results in the evolutionary optimization of its limited resources. A finely-tuned regulation allows rapid and efficient changes in protein levels [17], such as the repression of protein synthesis during cellular stress and enhancement of stress-responsive protein translation [18,19]. Together with the activity *in trans* of miRNAs and potentially of SINEUPs, this is mainly achieved through regulatory RNA sequences acting in *cis* during translation initiation [17,20], such as Internal Ribosome Entry Sites (IRES) [19,21] and upstream open reading frames (uORFs) [22].

IRESs were first discovered as complex structures in the 5’UTRs of picornavirus transcripts and were later found to occur in other viral and cellular mRNAs [23]. IRESs can either interact with translation initiation factors or directly with the small ribosomal subunit leading to ribosomal positioning at or near the initiation codon and promoting translation initiation. Their activity is regulated by RNA-binding proteins (RBPs) known as IRES *trans*-acting factors (ITAFs) [24]. Recently, IRES sequences have also been found in several circular RNAs (circRNAs) [25–27]. circRNAs are single-stranded, covalently closed RNA molecules produced from pre-mRNAs through a process called backsplicing. With the advent of next-generation sequencing, thousands of circRNAs have been identified. Compared to their linear counterparts, they are independently controlled and are mostly conserved [28–30]. circRNAs can regulate gene expression at both transcriptional level, by associating to chromatin around the promoter of their cognate linear gene, or post transcriptionally by sponging miRNAs or RBPs, thus participating in the stress response [31]. Importantly, hundreds of circRNAs present embedded IRES sequences that can promote coding sequence (CDS) translation, representing a new class of protein-coding circularized RNAs [25,27].

In this study, we aim to demonstrate that the invSINEB2 sequence of *AS Uchl1* is an IRES when acting in *cis*, that known IRES sequences can act as Effector Domain in synthetic SINEUPs and that natural IRES-containing SINEUPs can enhance the translation of target mRNAs in circularized conformation.

## Results

### The invSINEB2 sequence of *AS Uchl1* acts as an IRES in *cis*

The natural SINEUP *AS Uchl1* RNA upregulates UCHL1 protein expression post-transcriptionally [2,3] (Figure 1a). This activity requires both the invSINEB2 sequence (ED) and the overlapping antisense region (BD). Synthetic *SINEUP-DJ-1* was designed from the *AS Uchl1* RNA, replacing the BD sequence with one targeting *DJ-1* mRNA from −40 to +4 nt, with +1 being the A of the translation start site AUG (Figure 1a) [3]. This artificial RNA sequence increased endogenous DJ-1 protein expression with no effects on its mRNA levels (Figures 1b and S1a,b). When the invSINEB2 (*SINEUP ΔED*) and the BD sequence (*SINEUP ΔBD*) were independently deleted, no activity was observed (Figure 1b). To assess the interaction between SINEUP lncRNA and its target mRNA, we performed a pull-down experiment with 1% paraformaldehyde (PFA) as a cross-linker and a 115 nts biotinylated RNA probe targeting *SINEUP-DJ-1*. Endogenous *DJ-1* mRNA was significantly enriched compared to the control upon *SINEUP-DJ-1* RNA pull-down, proving lncRNA/mRNA interaction *in vivo* (Figure 1c). As expected, *DJ-1* mRNA levels did not change upon *SINEUP-DJ-1* expression (Figure S1c,d).

To shed light on the molecular mechanism of SINEUP RNA activity, we applied polysome fractionation to HEK293T cells overexpressing *SINEUP-DJ-1* and showing an increased expression of DJ-1 protein. The total lysate was layered on a 15-50% sucrose gradient column and separated into 14 fractions (Figure 1d).

The analysis revealed that *SINEUP-DJ-1* RNA is highly enriched in the 40S/60S/80S fractions, while it is depleted from polysome-associated ones (Figure 1e), in line with previous studies [32]. The overall SINEUP distribution in the 40S/60S containing fractions was maintained in the absence of its binding domain (*SINEUP-ΔBD*) suggesting that the invSINEB2 sequence is important for co-sedimenting with 40S/60S ribosomal subunits. Interestingly, the BD deletion significantly decreased the sedimentation of SINEUP RNA with the 80S-containing fraction (Figure 1e). As expected, upon *SINEUP-DJ-1* overexpression, a statistically significant increase, from 15% to 22%, in *DJ-1* mRNA association with polysomes was observed (Figure 1f). By contrast, in accordance with previous experiments [3–5], total RNA showed no variations of endogenous total *DJ-1* mRNA levels (Figure S1e,f). *GAPDH* mRNA was used as a reference (Figure S1g).

Since natural *AS Uchl1* activity is triggered by stress-signaling pathways [2] and SINEUP RNA co-sediments with ribosomal subunits-containing fractions, we investigated whether the invSINEB2 element could act as an IRES sequence in *cis*. To this end, we first took advantage of a bicistronic activity assay where dual-luciferase reporter pRUF plasmids were used to compare the expression of Rluc and Fluc upon transient transfection in HEK293T cells (Figure 2). To assess IRES activity, the *AS Uchl1* invSINEB2 sequence was cloned in inverted or direct orientation between the two CDSs. *c-myc* IRES sequence was used as positive control and empty pRUF as a negative one (Figure 2a) [33,34]. While Rluc activity was found similar among all 4 constructs (Figure S2a), Fluc activity significantly increased in the presence of the SINEB2 sequence in inverted orientation or of the *c-myc*-IRES (Figure 2b). SINEB2, in direct orientation, did not show IRES-like activity (Figure 2b) mirroring previous results showing that it does not support the SINEUP activity of *AS Uchl1* [2]. The IRES-like activity of the invSINEB2 element was also observed in U2OS cells (Figure S2a). Given that the experimental definition of IRES sequences is prone to false positives due to cryptic promoters and/or splicing, several additional control experiments were carried out in HEK293T cells. Firstly, the IRES activity of invSINEB2 was assessed by over-expressing a promoter-less pRUF plasmid. Despite the absence of the SV40 promoter, a low amount of bicistronic RNA was transcribed (Figure S2b-h) in accordance with previous studies showing the presence of cryptic promoter activity independent from tested sequences [35]. Nevertheless, both Rluc and Fluc activity strongly decreased upon SV40 promoter deletion, proving that IRES activity *in cis* is not due to the invSINEB2 sequence acting as a cryptic promoter (Figure S2b-h). These results were confirmed by looking at the RNA levels of both Renilla and Firefly CDSs that also excluded the presence of potential alternative splicing events (Figure S2i-n). In addition, bicistronic RNA was *in vitro* transcribed, capped and polyadenylated and directly transfected in equal amounts in HEK293T cells. As shown in Figure 2c, the IRES activity of the invSINEB2 sequence was similar to the one of *c-myc IRES*, the positive control, in conditions where the expression and the integrity of the IVT mRNA were experimentally monitored (Figure S2o-t). Last but not least, we took advantage of a vector system producing translatable circular mRNAs as a proof of concept that invSINEB2 acts in a cap-independent fashion. Briefly, the vector contains a single exon encoding two GFP fragments in a reversed order, and the IRES sequence is inserted upstream of the GFP start codon to drive cap-independent protein synthesis upon backsplicing (Figure 2d). The backsplicing is generated by the presence of two complementary introns that flank the GFP exon fragments which bring splice sites in proximity to produce a circRNA coding for GFP protein (Figure 2d). As a negative control, a GFP-coding circRNA with an inactive IRES sequence was used. When embedded in the circular RNA, the invSINEB2 drove the synthesis of GFP protein in a cap-independent manner (Figure 2e). Circular and linear RNA levels were measured by qRT-PCR analysis (Figure S2u-v).

**Figure 2.**
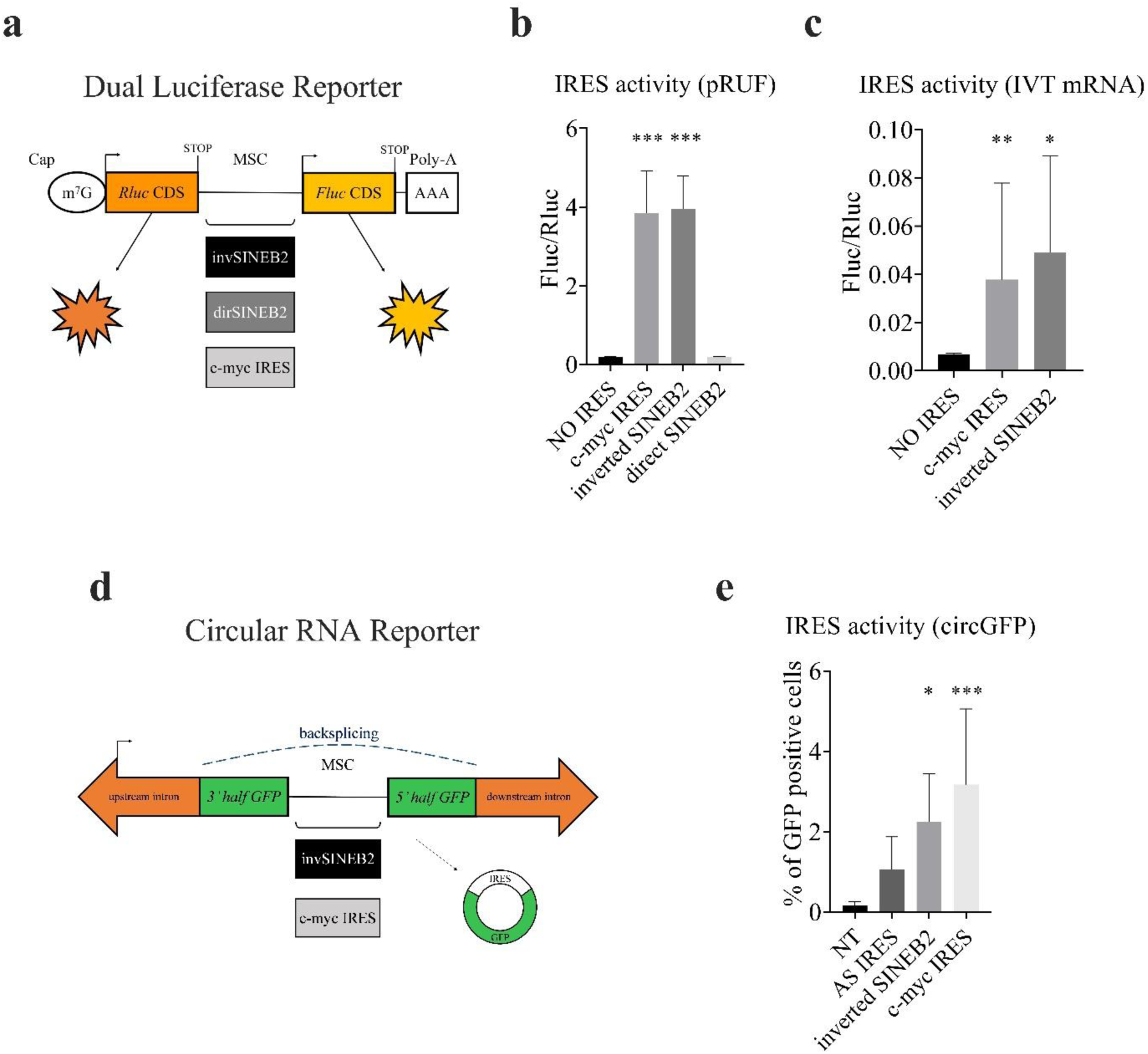
The invSINEB2 sequence presents IRES activity *in cis*. **a)** Design of pRUF-dual luciferase reporter vectors. Empty pRUF-MCS comprises a sequence codifying for Renilla luciferase (Rluc, orange) and a CDS for the Firefly luciferase (Fluc, yellow). Rluc CDS is constitutively expressed and translated in a cap-dependent manner; an MCS is placed between Rluc and Fluc CDSs. Fluc expression depends on the presence of a candidate upstream sequence which drives protein translation. SINEB2 element (from *UChl1* RNA) was cloned in direct (dirSINEB2, grey) or inverted (invSINEB2, black) orientation between Rluc and Fluc sequences, as well as *c-myc* IRES (light grey). **b)** IRES activity. Empty pRUF-MCS (with NO IRES between the two CDSs) and pRUF-*c-myc* IRES were used as negative and positive controls respectively. Luciferase activity was measured 48 h after transfection. Histogram shows that the SINEB2 element, in inverted orientation, induced Fluc protein translation in *cis*. Results are shown as the detected activity ratio of Fluc and Rluc (Fluc/Rluc) normalized against the empty pRUF-MCS background signal. IRES activity was calculated compared to the negative control. Plot indicates mean ± SD of 4 biological replicates performed in duplicate; ***p <0.001 were considered significant. Ordinary one-way ANOVA (with Holm-Sidak’s multiple comparisons test) was used to calculate significances. **c)** IRES activity (IVT mRNA). Bicistronic mRNAs were *in vitro* transcribed and transfected in HEK293T cells. Bicistronic mRNA without an IRES sequence between the two CDSs (NO IRES) and bicistronic mRNA with *c-myc* IRES were used as negative and positive controls respectively. Luciferase activities were measured 6 h following transfection. Histogram shows that the SINEB2 element, in inverted orientation, induced Fluc protein translation in *cis*. Results are shown as the detected activity ratio of Fluc and Rluc (Fluc/Rluc) calculated compared to the negative control (NO IRES). Plot indicates mean ± SD of 4 biological replicates performed in duplicate; **p <0.01, ***p <0.001 were considered significant. One-way ANOVA was used to calculate significances. **d)** Design of circular RNA reporter vectors. pCDNA3.1 (+) ZKSCAN 1 MCS-WT Split GFP + SENSE IRES is composed of two complementary introns flanking two GFP exon fragments (in reversed order) to produce a circRNA encoding GFP protein upon backsplicing. GFP expression depends on the presence of an upstream sequence which drives a cap-independent protein translation. SINEB2 element (from AS *Uchl1* RNA) was cloned in inverted (invSINEB2, black) orientation between the two GFP fragments, as well as *c-myc* IRES (light grey). **e)** IRES activity (circGFP RNA). circGFP RNAs were transfected in HEK293T cells. Untransfected cells and circGFP RNA with an IRES in inverted orientation between the two GFP coding fragments were used as negative controls. circGFP RNA with *c-myc* IRES was the positive control. Fluorescence was measured at 48 h following transfection through a cytofluorimeter. Histogram shows that SINEB2 element, in inverted orientation, induced GFP protein translation in a cap-independent manner. Results show the fluorescence percentage. Plot indicates mean ± SD of 5 biological replicates; **p <0.01, ***p <0.001 were considered significant. Ordinary one-way ANOVA was used to calculate significances.

In summary, we found that SINEUP RNA co-sediments with the 40S/60S/80S ribosomal subunits-containing fractions and that the SINEB2 element exerts IRES activity in *cis* when present in inverted orientation. These results provide an important information on the molecular mechanism of SINEUP activity and define a previously undisclosed biological function of embedded TEs.

### Viral and cellular IRES sequences function as ED in synthetic SINEUP RNAs

These data intriguingly suggest that natural IRES sequences could act as ED when embedded in a synthetic SINEUP RNA. To test this hypothesis, a synthetic *SINEUP(c-myc IRES)-DJ-1* RNA, with the *c-myc* IRES sequence substituting the invSINEB2, was transiently expressed in HEK293T cells (Figure 3a). *SINEUP-DJ-1* was used as a positive control. As reported in Figure 3b, both *SINEUP-DJ-1* and *SINEUP(c-myc IRES)-DJ-1* caused a reproducible increase in DJ-1 protein levels (1.8 fold) compared to the control. No changes in *DJ-1* mRNA levels were observed with qRT-PCR analysis, confirming SINEUP RNAs act post-transcriptionally (Figure S3a,b). These results were not cell specific as they were confirmed in HepG2 cells (Figure S3c-e).

**Figure 3.**
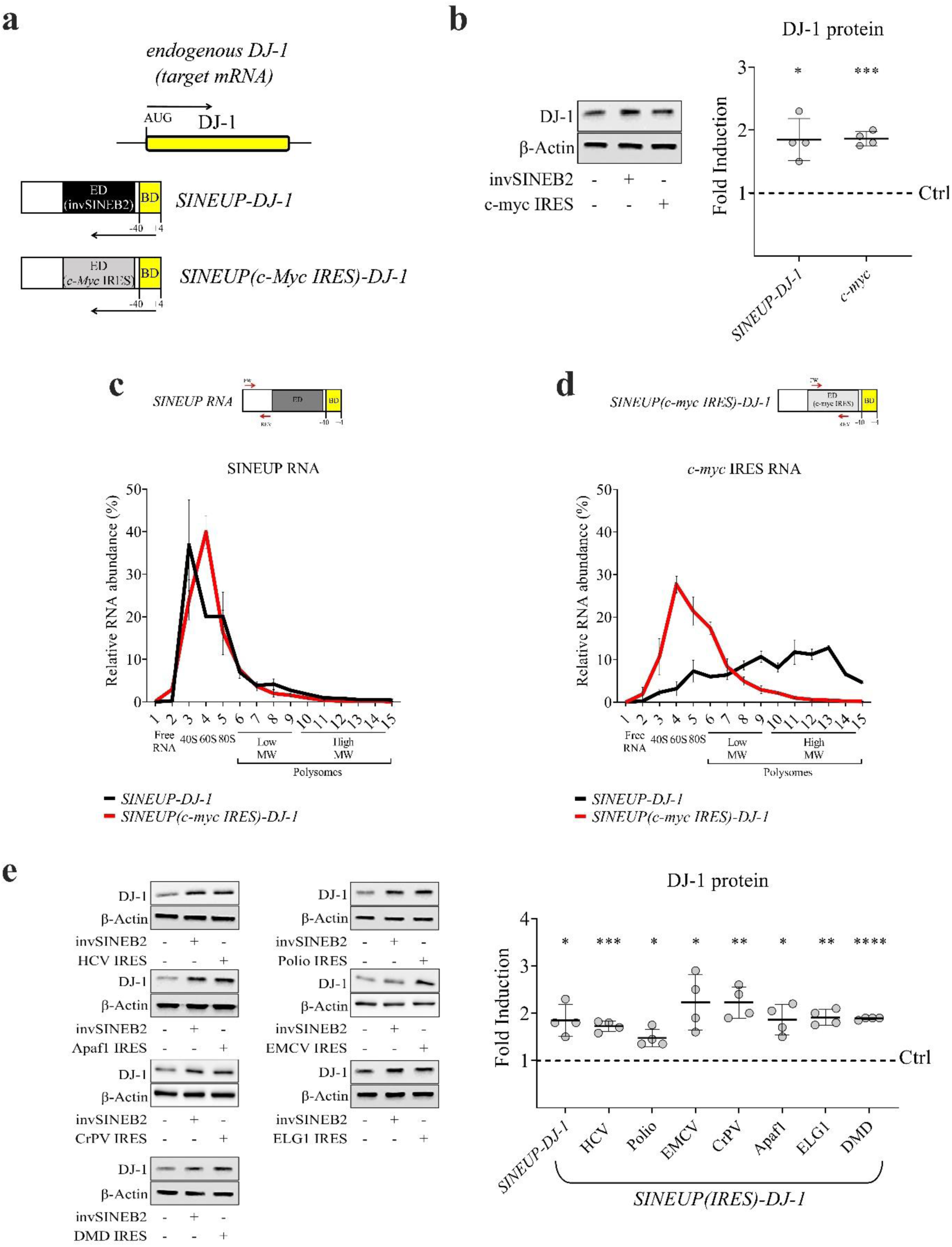
Cellular and viral IRESs increase DJ-1 protein synthesis acting as ED in SINEUP. **a)** Schematic overview of *SINEUP-DJ-1* and *SINEUP(c-myc IRES)-DJ-1*. Canonical synthetic SINEUP targeting *DJ-1* mRNA (yellow) comprising a BD (yellow) and an invSINEB2 as ED (black). *SINEUP(c-myc IRES)-DJ-1* comprises a BD, targeting *DJ-1* mRNA (−40/+4) in an antisense orientation, and the *c-myc* IRES sequence acting as potential EDs (light grey). **b)** Representative western blot image and reassuming plot reporting *SINEUP-DJ-1* and *SINEUP(c-myc IRES)-DJ-1* activity. SINEUP constructs were transiently expressed in HEK 293T cells and their activity was measured 48 h after transfection with western blot using an anti-DJ-1 antibody to detect and quantify DJ-1 protein. β-actin was used to normalize. Empty pCS2^+^ was used as control (Ctrl=1, dotted line). SINEUP RNAs increased DJ-1 protein levels. Plot reports DJ-1 fold induction mean ± SD; it represents 4 independent biological replicates performed in duplicate; *p <0.05, **p <0.01, ***p <0.001 and ****p <0.0001 were analyzed with ordinary one-way ANOVA (Holm-Sidak’s multiple comparisons test). **c)** SINEUP RNA enrichment. The relative abundances (in %) of *SINEUP-DJ-1* and *SINEUP(c-myc IRES)-DJ-1* were analyzed by RT-qPCR in each of the 14 gradient fractions. RNA distribution was reported as the mean ± SD of 3 replicates. **d)** *IRES* distribution as ED in SINEUP RNA and endogenous *c-myc* mRNA. The enrichment (in %) of *c-myc* IRES was analyzed, in the presence of *SINEUP(c-myc)-DJ-1* or of the canonical *SINEUP-DJ-1*, by RT-qPCR in each of the 14 gradient fractions. RNA distributions were reported as the mean ± SD of 3 replicates. **e)** Representative western blot images and reassuming plot reporting *SINEUP(IRES)-DJ-1* activity. SINEUP constructs were transfected in HEK293T cells and their activity was measured 48 h later with western blot. Western blot analysis was carried out using anti-DJ-1 antibody to detect and quantify DJ-1 protein. β-actin was used to normalize. Empty pCS2^+^ was used as control (Ctrl=1, dotted line). All SINEUP(IRES) RNAs increased DJ-1 protein levels. Plot reports DJ-1 fold induction mean ± SD; it represents 4 independent biological replicates performed in duplicate; *p <0.05, **p <0.01, ***p <0.001 and ****p <0.0001 were analyzed with ordinary one-way ANOVA (Holm-Sidak’s multiple comparisons test). All SINEUP(IRES)s were designed using a natural SINEUP backbone, derived from Δ5’-*AS Uchl1*, replacing the SINEB2 sequence with IRES elements of different origins.

To study the molecular mechanism by which SINEUP(IRES) can enhance translation, we applied polysome fractionation to HEK293T cells overexpressing *SINEUP(c-myc IRES)-DJ-1*. The analysis revealed that *SINEUP(c-myc IRES)-DJ-1* RNA is highly enriched in the 40S/60S/80S fractions, while it is depleted from polysome-associated ones, as in the case of *SINEUP-DJ-1* (Figure 3c). On the contrary, the endogenous *IRES* sequence, embedded in *c-myc* mRNA, co-sedimented with the 40S/60S, 80S monosomes and polysomes fractions, as expected for a protein coding transcript (Figure 3d). This pattern was confirmed when primers were amplifying the 3’UTR region of the endogenous *c-myc* mRNA (Figure S3f). Taken together, these results state that *c-myc IRES* shows a different, atypical distribution upon polysome profiling when embedded in a synthetic SINEUP RNA.

Since viral sequences are classified into four types according to their mechanisms of action and their requirement of specific translation factors, a representative example for each IRES class was then assayed: poliovirus (Polio) (type I), encephalomyocarditis virus (EMCV) (type II), hepatitis C virus (HCV) (type III) and cricket paralysis virus (CrPV) (type IV) [36]. Concerning cellular IRES sequences, we investigated the activity of the following IRES-containing mRNAs: apoptotic peptidase activating factor 1 (*Apaf1*), enhanced level of genomic instability 1 homolog (*ELG1*) and dystrophin (*DMD*). Accession numbers and RNA sequences used in these experiments are indicated in Table S1 (http://www.iresite.org). The canonical synthetic *SINEUP-DJ-1* was used as a positive control. As reported in Figure 3e, transient expression of *SINEUP(IRES)-DJ-1* RNAs in HEK293T cells caused a significant increase in DJ-1 protein levels similar or higher than the positive control, with *SINEUP(EMCV IRES)-DJ-1* and *SINEUP(CrPV IRES)-DJ-1* reaching the highest activity levels (>= 2-fold). No changes in *DJ-1* mRNA were observed with qRT-PCR analysis, confirming a post-transcriptional control (Figure S3a,b). Activity was confirmed in HepG2 cells for all IRES sequences, with some differences in the amount of fold changes (Figure S3c-e). Given that the combination of simply ED and BD sequences (miniSINEUP) recapitulates the ability of full-length SINEUPs to increase target protein expression, we then examined whether constructs solely containing IRES sequences and DJ-1 BD (*miniSINEUP(IRES)-DJ-1s*) retained the same activity level of longer constructs (Figure S3g-j). Transient transfection in HEK293T cells led to a significant increase in DJ-1 endogenous protein levels (Figure S3h) with no changes in the amount of *DJ-1* mRNA (Figure S3i).

Taken together, these results demonstrate that established viral and cellular IRES RNAs can act as ED in synthetic SINEUPs increasing protein expression of a target through the presence of an antisense sequence.

### *SINEUP(IRES)* RNA maintains its activity in circularized conformation

Building on the observations that the invSINEB2 of *AS Uchl1* RNA functions as an IRES in *cis* and that viral and cellular IRES RNAs can act as ED in synthetic SINEUPs, we aimed to identify endogenous non-coding RNAs with an embedded IRES sequence and exhibiting SINEUP-like activity.

In contrast with the invSINEB2 of natural SINEUPs, IRES-containing RNAs mostly correspond to protein-coding transcripts. Furthermore, target mRNAs cannot be identified for their localization within a sense/antisense pair in the genome. Therefore, we decided to focus on a specific RNA class, circRNAs, arising from protein-coding genes and reported to include IRES RNA sequences. circRNAs can be both non-coding and coding for proteins [25,37,38]. In the latter case, IRES sequences promote translation in *cis* of CDS included in the circularized RNAs [39].

As a proof-of-principle that circularized *SINEUP(IRES)* RNAs are post-transcriptionally active as their linear counterparts, we generated a synthetic circularized SINEUP RNA (circSINEUP) combining *c-myc* IRES element as ED and a previously established GFP targeting sequence as BD, with the circularizing pcDNA3.1(+) ZKSCAN1 MCS Exon Vector (Figure 4a). The sequence was inserted between two highly complementary intronic repeats (derived from the *zkscan* gene) that allow the circularization of the inserted sequence [40]. An empty pcDNA3.1(+) ZKSCAN1 MCS Exon Vector was used as a negative control. This construct was named *circSINEUP(c-myc IRES)-GFP*. Linear *miniSINEUP-GFP* [41] and *miniSINEUP(c-myc IRES)-GFP* were used as positive controls (Figure 4). Upon co-transfection of these constructs with one expressing GFP (pEGFP) in HEK293T cells, *circSINEUP(c-myc IRES)-GFP* induced an increase in GFP protein expression of 2.2 fold, similar to canonical linear SINEUPs (Figure 4b). Transcripts’ circularization was confirmed with divergent primers amplification (data not shown). As expected, no variation in GFP mRNA levels was observed, confirming that both linear and circularized SINEUPs act post-transcriptionally (Figure S4). These results demonstrate that synthetically designed circSINEUPs retain SINEUP activity.

**Figure 4.**
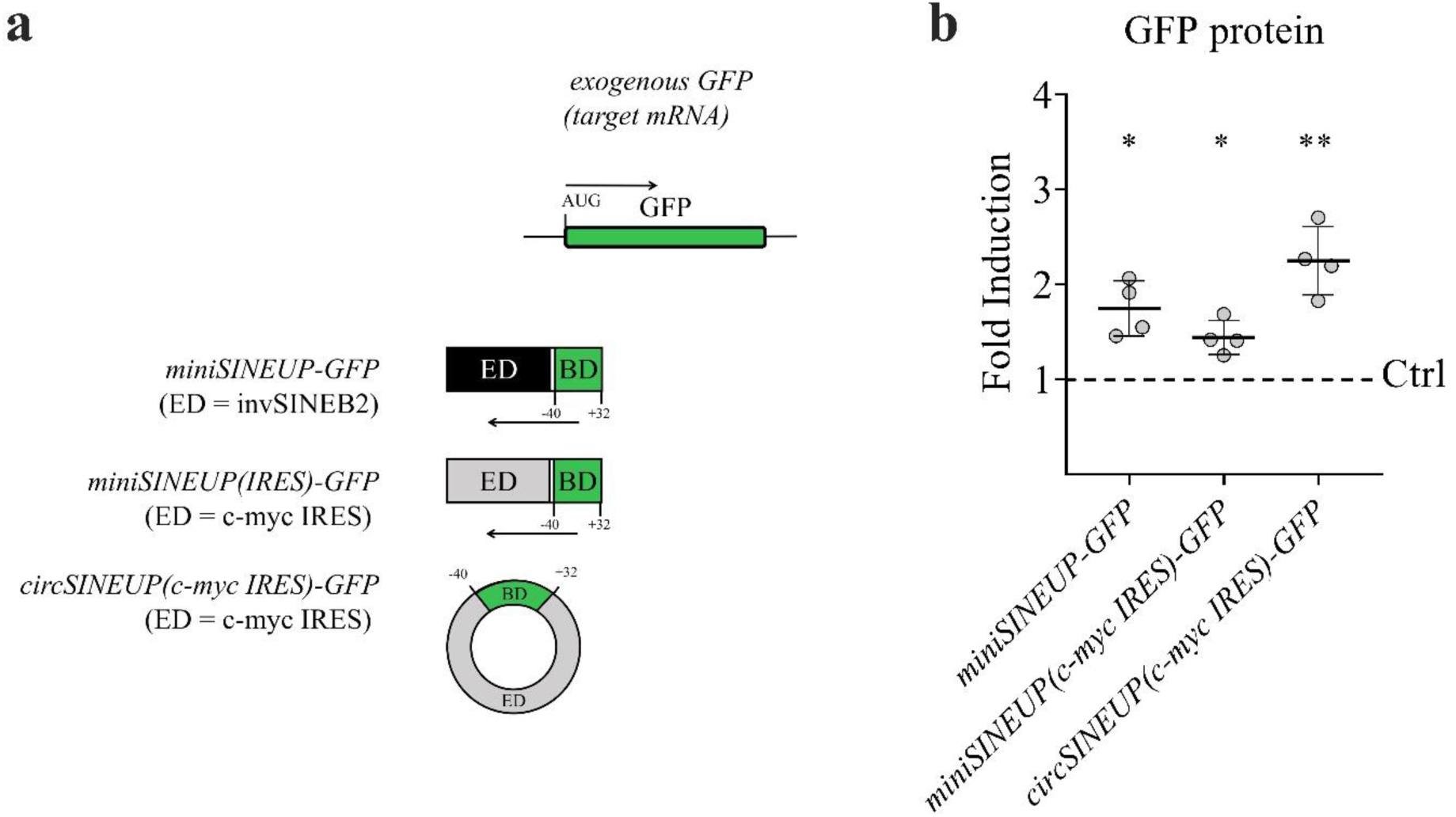
Circularized SINEUP(IRES) RNA maintains its activity in *trans*. a) Schematic representation of circularized SINEUP(IRES) RNA (circSINEUP). A synthetic *SINEUP(c-myc IRES)-GFP* comprising a BD (green) targeting *GFP* mRNA (−40/+4, green) and a *c-myc* IRES element as ED (light grey) was cloned into pcDNA3.1(+)ZKSCAN MCS Exon vector to be circularized. **b)** *circSINEUP(c-myc IRES)-GFP* activity. HEK293T cells were co-transfected with a circRNA-expressing vector and an EGFP-expressing one in a 6:1 ratio. 48 h after transfection, GFP protein fluorescence was measured. GFP fluorescence intensity was normalized on the NucBlue signal. Empty pCDNA3.1(+) was used as the negative control (Ctrl=1, dotted line) and linear *miniSINEUP-GFP* and *miniSINEUP(c-myc IRES)-GFP* were the positive controls. *circSINEUP(c-myc IRES)-GFP* increased GFP protein levels more than the linear positive controls. The dot plot reports GFP fold induction and the mean ± SD from 4 independent biological replicates; *p <0.05, **p <0.01 and ****p <0.0001 were calculated with ordinary one-way ANOVA (Holm-Sidak’s multiple comparisons test).

### *circ5533* is a natural circSINEUP RNA that enhances *Pxk* translation

To investigate the existence of natural circSINEUPs, we took advantage of the circBase database and of a list of sequences present in cellular and viral RNAs reported to support IRES activity in bicistronic assays [42,43]. In total, 140,790 unique IDs from circBase were mapped on the human genome, from which 570 sequences with IRES activity were retrieved from [42]. The 570 trimmed IRES sequences, of a length of 174 nts, were aligned using the BLAST tool against the 140,790 circRNA sequences. Considering only the alignments reaching 100% IRES coverage and mismatches <=1 nt without gaps, 411 unique circRNAs containing sequences with potential IRES activity were identified (Table S2). We then filtered out circRNAs with ORF sequences longer than 100 aa to look for potential non-coding circRNAs. The Virtual Ribosome tool (Dna2pep v1.1)[44] was used to predict the longest complete ORF for each circRNA by searching across all positive reading frames with methionine as the start codon and a canonical stop. 49 circRNAs from 32 genomic loci presented a predicted ORF of 100 or less aa in length (Table 1). Among them, only hsa_circ_0085533 (*circ5533)* showed the absence of ORF according to the used selection criteria. *circ5533* is a circular non-coding RNA transcribed from the *c-myc locus* that is 555-nts-long and contains an IRES sequence of 380 nts. This is usually embedded in the 5’UTR of *c-myc* linear mRNA, with well-established evidence of activity in *cis* [45]. In addition, as reported above, *c-myc* IRES can act as ED in synthetic linear SINEUP (Figure 3 a,b, and Figure 4) and circSINEUP (Figure 4). Notably, *circ5533* also presents 175 nts of unknown function that could act as BD sequences (Figure 5a). To investigate *circ5533*’s ability to function as a circular SINEUP and identify its potential target mRNAs, we overexpressed *circ5533* RNA in HEK293T cells, taking advantage of the circularizing vector previously used for *circSINEUP(c-myc IRES)-GFP* expression. Changes in proteins expression were then analyzed through untargeted proteomics and coupled with RNA-seq to identify proteins with increased expression and no change in the respective mRNA level. Experiments were carried out in six biological replicates. With the proteomic analysis, 117 proteins, whose RNA expression level were not affected (Figure S5a), were found significantly upregulated in *circ5533*-transfected cells compared to controls (Figure 5b). Notably, the absence of significant changes in mRNA levels strongly suggests there was no potential sponge effect on miRNA activity in our experimental settings. As a validation, we chose six representative examples from the most reproducibly increased proteins that could be efficiently detected with commercially available antibodies. Western blot analysis confirmed statistically significant increase of four out of six proteins tested (PXK, MFN, CHK1, GUCY1B1) (Figure S5b). RT-qPCR analysis of corresponding transcripts confirmed that mRNA levels were not changed in cells overexpressing *circ5533* compared to control (Figure S5c,d). We focused on *Pxk* as a representative mRNA target of circSINEUP activity mediated by *circ5533*. To rule out any possibility that *circ5533* activity was dependent on the type of circularizing plasmid, we cloned *circ5533* annotated sequence (NM_002467) into a second circularizing vector named pcDNA3.1(+) Laccase2 MCS Exon Vector, and demonstrated that *circ5533* activity was able to increase endogenous PXK protein levels to 1.7 fold with no change in mRNA levels (Figure 5c and Figure S5e,f). To assess whether *circ5533* was acting at the translational level, we carried out polysome fractionation analysis of *circ5533*-overexpressing HEK293T cells. An empty vector was transfected as a negative control. *circ5533* RNA was significantly enriched mainly in the fractions containing 80S monosomes and polysomes, suggesting its interaction with the translation machinery (Figure 5d). Upon *circ5533* overexpression, a consistent and significant increase of *Pxk* mRNA co-sedimentation with polysomes was observed (Figure 5d). Total *Pxk* mRNA levels did not change in total RNA extracts (Figure S5j,k). *GAPDH* mRNA distribution was used as <u>a</u> control to assess the specificity of the *circ5533* effect and as a reference for the accuracy of polysome fractionation analysis (Figure S5l).

**Figure 5.**
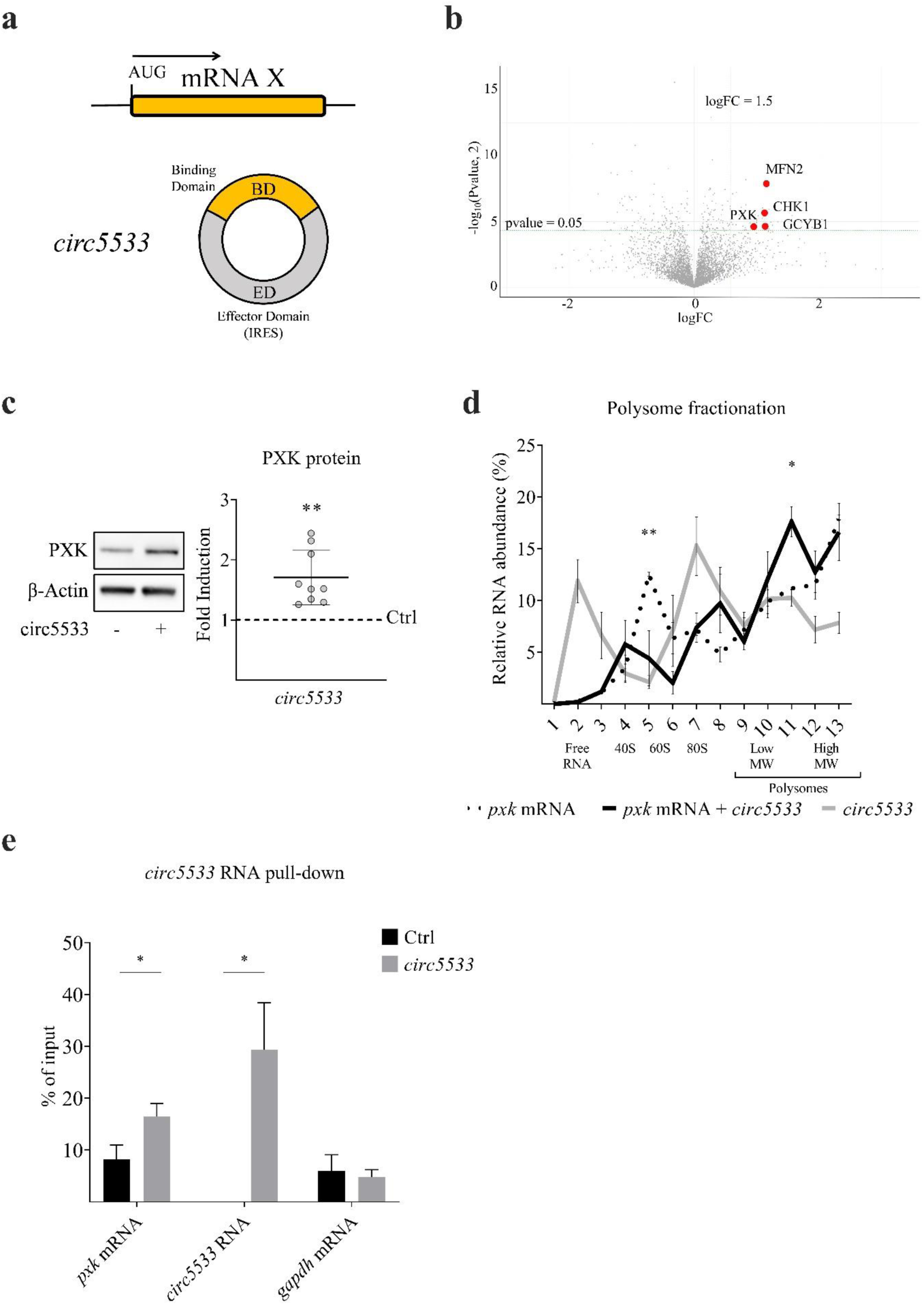
*Circ5533* is an IRES containing circRNA transcribed from the *c-myc* locus and it affects cellular proteome upon overexpression. **a)** Schematic representation of *circ5533*’s potential mechanism of action. *circ5533* comprises an IRES sequence (grey) as ED and an upstream region potentially acting as mono- or multi-BD (dark yellow) to one or more endogenous target mRNAs. **b)** *circ5533* acts as a circSINEUP increasing target protein levels. Experiments were performed as described in Methods (“Cell Lines and Transfection Conditions”). HEK293T cells were transfected with pcDNA3.1(+)-ZKSCAN-*circ5533* (n=6). Empty pcDNA3.1(+)-ZKSCAN was used as a negative control. Protein extract was analyzed through mass-spectrometry; resulting data were normalized assuming that the number of proteins, whose levels did not change, was larger than the number of proteins whose levels changed (MLR method). Fold-inductions were calculated compared to the negative control as reported in Methods (“Mass Spectrometry”). The volcano plot reports detected protein levels following *circ5533* overexpression. The log_2_ fold change (FC), on the x-axis indicates the log2 ratio between treated and control samples of the mean expression level of each protein; the – log_10_ p-values are plotted on the y-axis. Dots on the left, with respect to log FC=0 on the x-axis, represent downregulated proteins when *circ5533* is expressed. Dots on the right are proteins whose levels increased. Red dots represent proteins whose fold inductions (FC>1.5) were validated through a western blot. p <0.05 was considered significant. **c)** *circ5533* increases natural PXK protein levels. HEK293T cells were transfected with pcDNA3.1(+)- Laccase2-*circ5533* (n=9). Empty pcDNA3.1(+)-Laccase2 vector was the negative control. Western blot analysis was performed, Anti-PXK Ab was used to detect PXK protein. PXK fold induction was calculated compared to the empty control (Ctrl=1, dotted line). β-actin was used to normalize. The ability of *circ5533* to increase PXK was thus confirmed. *Circ5533* activity was plotted as mean fold induction values ± SD. **p <0.01 was considered significant (one-sample t-test). **d)** Lysates from control and treated cells were fractionated by sucrose gradients and separated through a UV detector continuously measuring the solution absorbance (260 nm). The relative distributions (in %) of *circ5533* RNA and *Pxk* mRNA were analyzed by RT-qPCR in each of the 13 gradient fractions. RNA distribution was reported as the mean ± SD of 3 replicates. *Pxk* mRNA increased in polysome fraction following *circ5533* over-expression. *p <0.05 and **p <0.01 were considered significant (2way ANOVA, Sidak’s multiple comparisons test). **e)** *circ5533* RNA Pull-Down (RPD). *circ5533* was purified using biotinylated DNA probes whose lengths range around 20 nts. As a negative control, RPD was performed in the absence of targeting probes. qRT-PCRs show the enrichment of *circ5533* RNA and *Pxk* mRNA upon *circ5533* RPD. Columns represent the mean enrichment over input (%) relative to the negative control ± SD from 6 independent biological replicates. Significant *p <0.05 values were analyzed with unpaired t-test.

**Tab 1.**
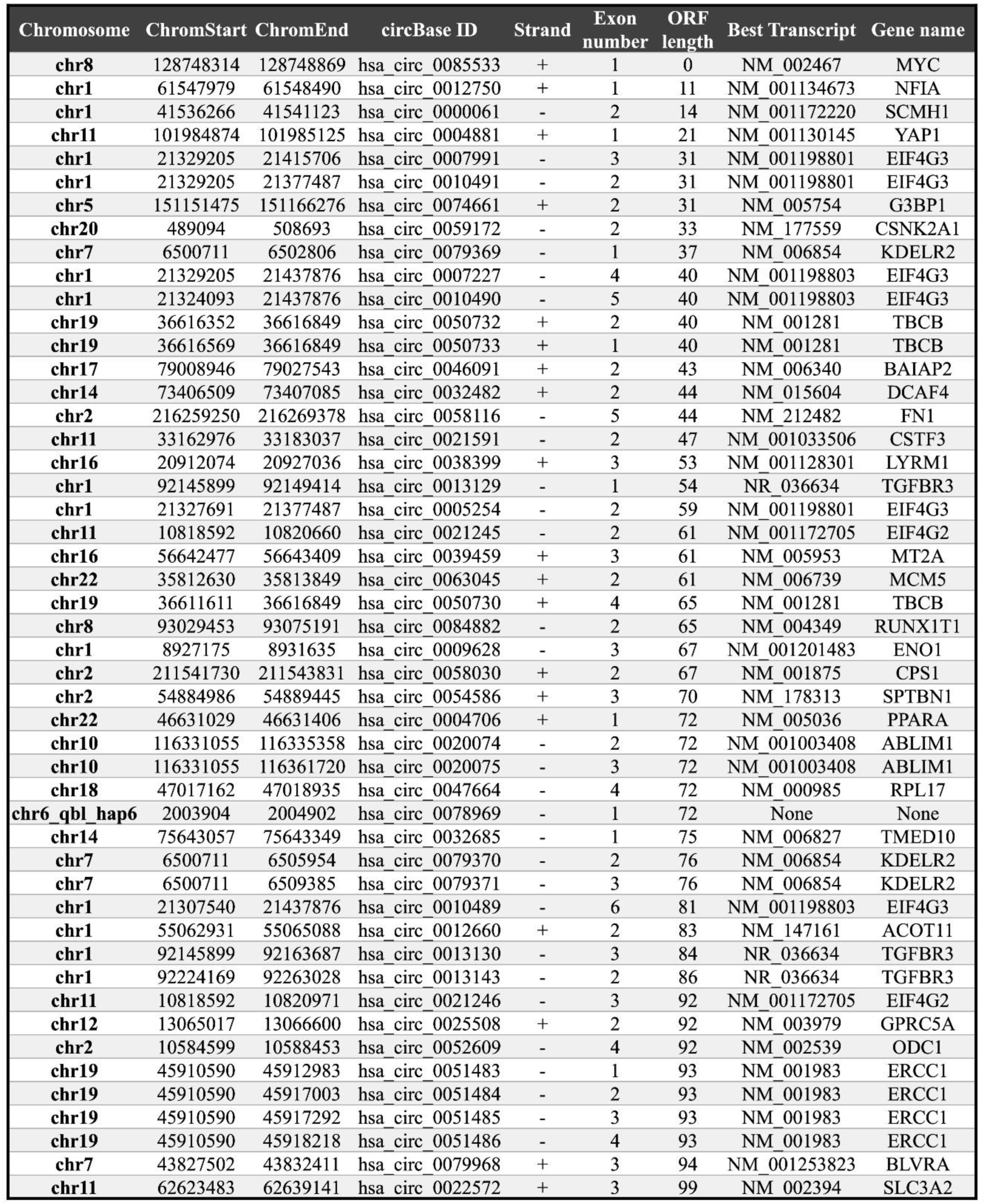
List of circRNAs containing an IRES sequence with a predicted ORF of 100 or less aa in length.

As demonstrated above for *SINEUP-DJ-1* and *DJ-1* mRNA, SINEUP activity requires a physical interaction between noncoding RNA and the mRNA target. To prove the formation of *circ5533* RNA and *Pxk* mRNA heteroduplexes, we performed a pull-down experiment with 1% paraformaldehyde (PFA) as cross-linker and a pool of 3 biotinylated DNA probes targeting *circ5533* (Figure 5e and Figure S5m). As expected, endogenous *Pxk* mRNA was significantly enriched upon *circ5533* pull-down when compared to the empty control (Figure 5e).

Furthermore, we carried out a co-transfection analysis with full-length *Pxk* cDNA fused to an *in- frame* C-terminal FLAG tag (WT PXK-FLAG). Detecting WT PXK-FLAG protein with an anti- FLAG antibody in western blot analysis, we confirmed that *circ5533* increased the amount of the target protein acting post-transcriptionally (Figure 6b,c, Figure S6a,b).

**Figure 6.**
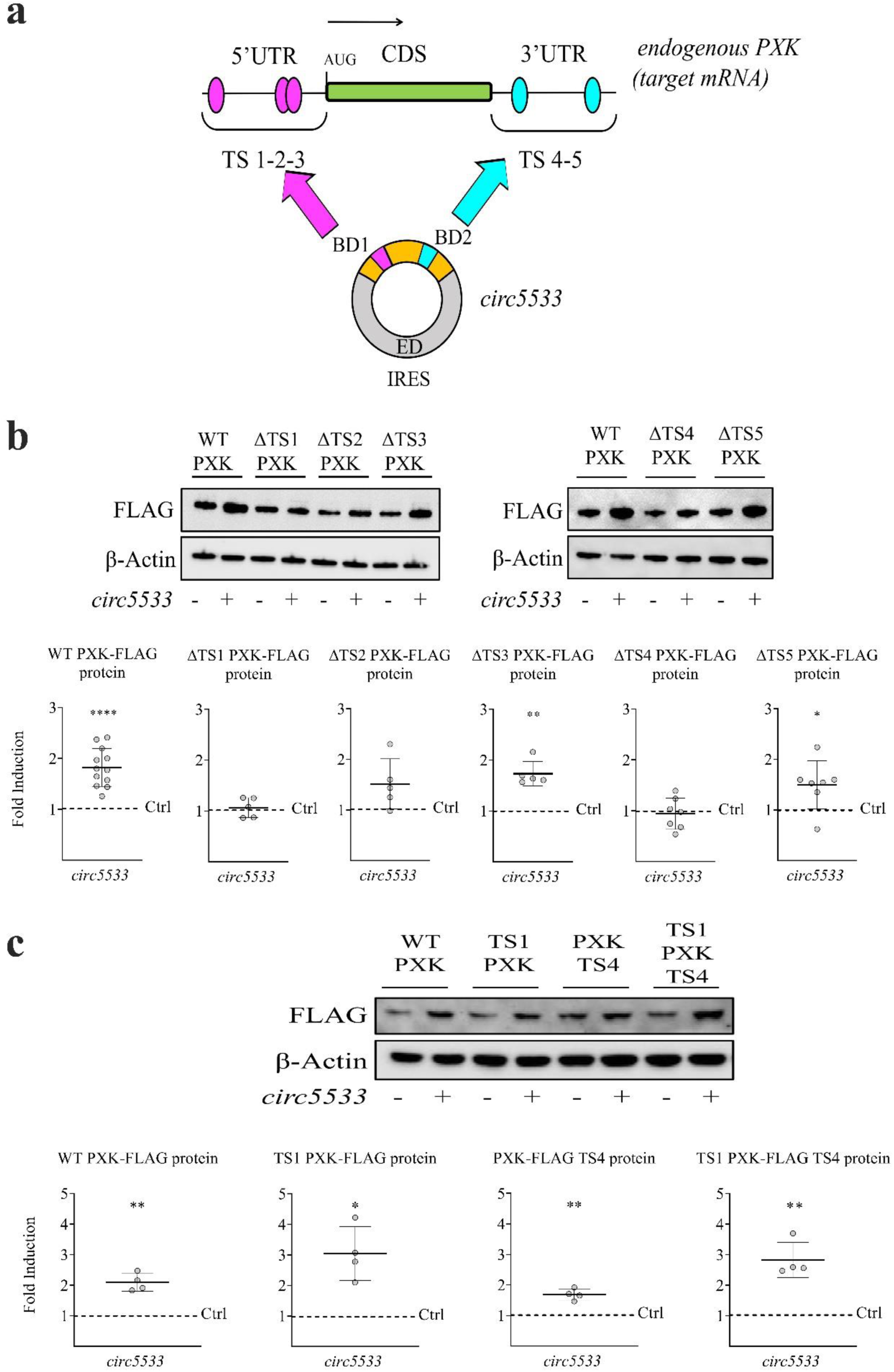
*circ5533* increases PXK protein expression through the activity of two BDs. **a)** Schematic design of putative Binding Domains (BDs) in *circ5533* RNA. Putative BD1 (pink) and BD2 (blue) are located on the region upstream (dark yellow) of the IRES sequence (grey) of *circ5533*. BDs were identified looking for the longest sequence pairing, in antisense orientation, between circ5533 RNA and *Pxk* mRNA; a minimal perfect match of 8nts was considered. BD1 can potentially target 3 different regions on the 5’UTR of *Pxk* mRNA, identified as target sites (TSs) 1, 2 and 3 (pink). BD2 could pair with TS4 and/or TS5 (blue) on the 3’UTR of *Pxk* mRNA. *Pxk* CDS is highlighted in green. **b)** *Circ5533* activity on PXK-FLAG mutants. Identified TSs were individually deleted from *Pxk-flag* RNA; each construct was co-transfected with *circ5533* in HEK293T cells (1:12). After 48 h, cells were harvested and proteins extracted. Empty pcDNA3.1(+)-Laccase2 vector, co-transfected with different *Pxk-flag* mutants, was used as a negative control; PXK-FLAG WT + *circ5533* was the positive control. Deletion of TS1 from *Pxk* 5’UTR or of TS4 from *Pxk* 3’UTR inhibited the *circ5533*-mediated increase in PXK-FLAG protein. Anti-FLAG Ab was used to detect ectopic PXK-FLAG. Fold inductions were calculated compared to the relative negative control (Ctrl=1, dotted line). β-actin was used to normalize. Plots report PXK-FLAG fold induction mean ± SD; they represent at least 5 independent biological replicates; *p <0.05, **p <0.01 and ****p <0.0001 were analyzed with one-sample t-test. **c)** Representative western blot image and reassuming plots of *circ5533* activity on PXK-FLAG mutants with only TS1 and/or TS4 at 5’ and 3’UTRs respectively. UTR regions were completely removed from *Pxk*-*flag* mRNA. Three new *Pxk-flag* mutated RNAs were assembled: TS1 PXK-FLAG with only TS1 at the 5’UTR; PXK-FLAG TS4 with only TS4 at the 3’UTR and TS1 PXK-FLAG TS4 with TS1 at the 5’ and TS4 at the 3’UTRs. Empty pcDNA3.1(+)- Laccase2 vector, co-transfected with different PXK-mutants, was used as a negative control; PXK-FLAG WT in co-transfection with *circ5533* was the positive control. TS1 on *Pxk* 5’UTR was sufficient to allow *circ5533*-mediated upregulation of PXK-FLAG protein. TS4 at the 3’UTR restored the protein fold induction at lower levels than the positive control. Anti-FLAG Ab was used to detect PXK-FLAG. Fold inductions were calculated compared to the relative negative control (Ctrl=1, dotted line). β-actin was used to normalize. Plots report PXK-FLAG fold induction mean ± SD from 4 independent biological replicates; *p <0.05 and **p <0.01 were calculated with one-sample t-test.

To rule out the miRNA-sponge activity of *circ5533*, we used a mouse embryonic stem cell (mESC) line specifically knocked-out for the DICER enzyme, a well-established experimental model to inhibit miRNA synthesis [46]. We co-transfected *circ5533* and the cDNA for *Pxk-flag* in both mESC WT and mESC *Dicer ^-/-^*. As shown in figure S5n-q, *circ5533* was able to increase PXK protein translation both in WT and *Dicer ^-/-^*, confirming that *circ5533* was not acting as a miRNA sponge.

To identify potential BDs in *circ5533* and target sites (TSs) in *Pxk* mRNA, we carried out a bioinformatic analysis looking for the longest pairing sequence in antisense orientation between these two transcripts. Matches with a minimal length of 8 nts between *Pxk* and the non-IRES sequence of *circ5533* were found. As described in Figure 6a and Table 2, three distinct potential TSs (TS1, TS2, TS3) in the 5’UTR of *Pxk* mRNA could pair to the same antisense region of *circ5533* from nucleotide 23 to 48 (named BD1). In addition, two potential TSs (TS4 and TS5) were identified in the 3’UTR of *Pxk* mRNA, pairing with a second antisense region (BD2) in the non-IRES sequence of *circ5533*, which spans from position 104 to 112. Interestingly, differently from linear natural SINEUP RNA, no TSs spanned the initiation AUG codon. We then deleted or inverted the orientation of both potential *circ5533* BD sequences to assess their effect on the activity. However, both mutations strongly diminished the expression level of *circ5533*, constraining this type of analysis (Figure S5g-i). We then proceeded by selectively mutating all five TSs on the *Pxk-flag* construct and verifying *circ5533* activity. As shown in Figure 6b and Figure S6a,b, the mutations on TS2, TS3, and TS5 did not alter *circ5533*’s effect, as detected with the anti-FLAG antibody. However, deletion of TS1 and TS4 inhibited the *circ5533*-mediated increase of PXK-FLAG protein, demonstrating that TS1 and TS4 are essential for *circ5533* activity and suggesting that both BD1 and BD2 are functionally active. We then investigated whether TS1 and TS4 were sufficient for establishing *circ5533* activity by designing three PXK-FLAG mutants: i. with only TS1 at the 5’UTR; ii. with only TS4 at the 3’UTR and iii. with both TS1 and TS4 at the 5’ and 3’UTRs respectively. Interestingly, the presence of TS1 alone was sufficient to fully restore *circ5533* activity, while the exclusive presence of TS4 increased protein expression to a lower extent. Importantly, the combination of TS1 and TS4 fully restored *circ5533* activity (Figure 6c). Efficient transcript circularization was assessed by amplifying with divergent primers (Figure S6d). No change in mRNA levels was observed in any of the experiments, confirming that the regulation occurs at the post-transcriptional level (Figure S6c).

**Tab 2.**
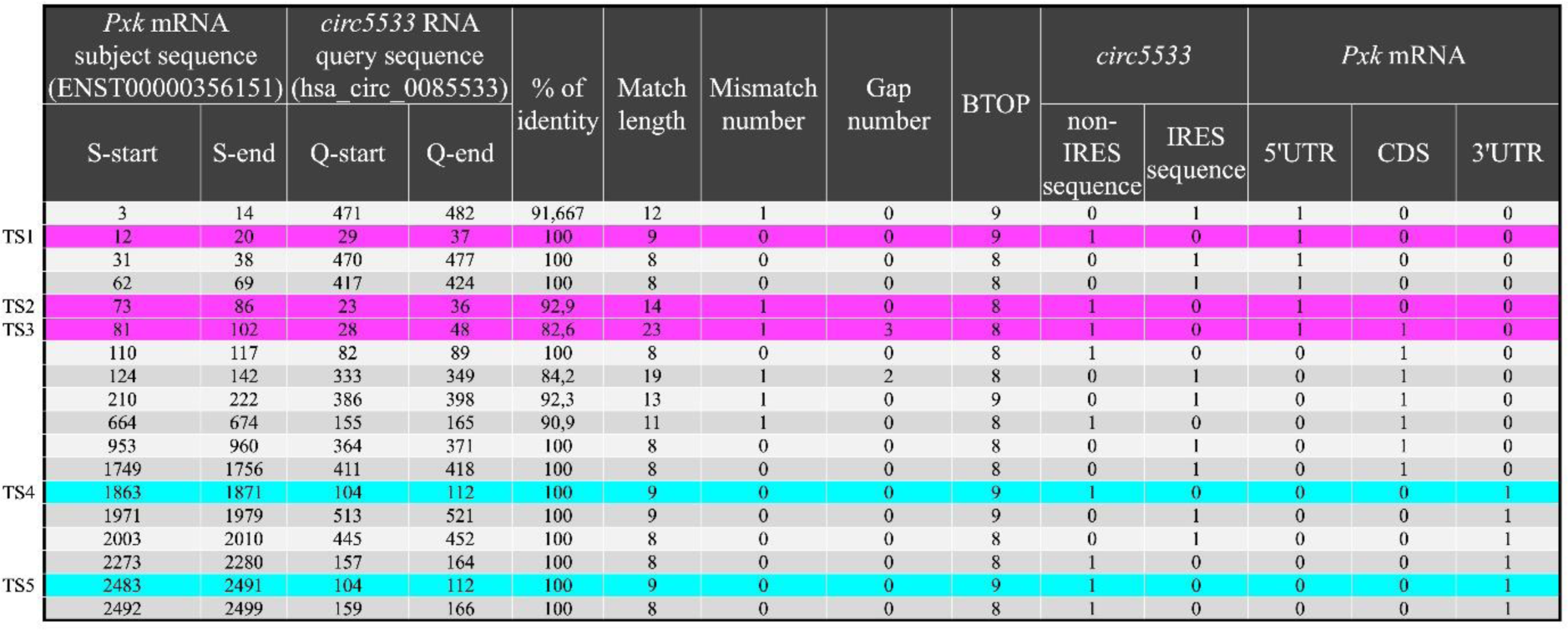
Identification of potentital BDs an TSs between *circ5533* and *<u>Pxk</u>* <u>mRNA.</u>.

Taken together, these results suggest a new functional role for circRNAs in modulating protein translation. We identified *circ5533* as a natural circSINEUP RNA able to increase PXK protein translation by interacting with its mRNA and the translational machinery.

## Discussion

The canonical definition of SINEUP is based on the combined activity of two RNA elements: a BD and an ED, where the ED is a SINEB2 sequence in an inverted configuration. Natural SINEUPs are part of genomic sense/antisense pairs in a head-to-head configuration [3]. Several examples are consistent with a model where they are regulatory RNAs that enhance the translation of specific mRNA targets during homeostatic response. This increase can be subtle but physiologically important given the tight regulation of protein levels *in vivo*, as shown by the pathological consequences of haploinsufficiency [47]. SINEUPs can also contribute to maintaining target mRNA translation in stress conditions when cap-dependent translation is attenuated [2,3].

SINEUP activity can therefore be interpreted as a cellular strategy to modulate translation of specific mRNAs when coping with stress. This is very similar to miRNAs function but with an opposite effect on protein levels, and to the activity of IRES sequences for their ability to drive translation of downstream CDS in *cis*. It is therefore intriguing that the invSINEB2 sequence from *AS Uchl1* is indeed able to act as an IRES in a series of assays for its activity in *cis*. This experimental observation increases the number of reported biochemical activities of embedded TEs in both coding and non-coding RNAs. It also strengthens the concept that RNAs are organized according to domains and extends the concept of the exaptation of TEs to RNA modules.

Polysome profiling shows that SINEUP RNAs co-sediment with the 40S/60S ribosomal subunits and 80S monosomes, while depleted from polysome fractions. This pattern depends on the invSINEB2 sequence, suggesting it contains all the elements required to interact with ribosomal subunits. By BD pairing, SINEUPs can interact with target mRNAs, enhance mRNAs’ association with heavy polysomes and increase endogenous levels of the encoded protein. It remains unclear whether the activity in *trans* of the invSINEB2 involves the same mechanistic details of an IRES sequence acting in *cis*.

Nevertheless, we showed that known viral and cellular IRES sequences share the ability to act as ED in synthetic SINEUPs. In the synthetic constructs tested in this study, IRESs act as ED in combination with an antisense sequence as BD, enhancing protein expression of an endogenous target mRNA. While IRESs are of different types and drive translation in *cis* with different mechanisms and requirements of cellular factors, it is unclear whether their abilities to act as ED in SINEUP RNAs are mechanistically similar to an IRES acting in *trans*. Furthermore, it is tempting to speculate that IRES sequences that belong to different types according to their activities in *cis* may present different mechanistic details when acting in *trans*. Further work is needed to address these important challenges. In this context, polysome profiling experiments showed that *c-myc* IRES sequences were behaving as a SINEUP RNA, being associated with the 40S/60S/80S-containing fractions and depleted from polysomes, when embedded as ED. As expected, they were associated with actively translating heavy polysomes when part of a canonical *c-myc* transcript.

IRES sequences are mostly present in protein-coding genes and our knowledge of their activity has been so far limited to their ability to promote translation in *cis*, usually under stress conditions. These results may imply that the very same IRES sequences can act as ED in SINEUP RNAs, affecting the cellular proteome in unpredicted ways. It is tempting to speculate that i.e. IRES-containing viral transcripts may affect the host cell proteome with a SINEUP(IRES)-like mechanism, potentially leading to the identification of cryptic BDs targeting cellular mRNAs in viral sequences. Protein-coding genes can indeed give rise to robust expression of uncharacterized IRES-containing lncRNAs [48]. To identify natural IRES-containing, non-coding RNAs with SINEUP-like activity, we focused on circRNAs since they represent a reservoir of transcribed IRES sequences and can be both coding and non-coding. First, we demonstrated that linear synthetic SINEUPs with *c-myc* IRES as ED can be active as circularized RNAs (circSINEUPs). Importantly, we listed a group of 49 non-coding circRNAs that present IRES sequences without a computationally predicted ORF longer than 100 aa, the conventional threshold for non-coding RNAs. It remains to be determined whether all these circRNAs are truly non-coding since some short polypeptides encoded by lncRNAs and circRNAs have been experimentally identified [37]. We then focused on *circ5533*, a circular RNA transcribed from the *c-myc* locus, as a potential natural circSINEUPs. *circ5533* was previously shown to be highly expressed in human melanoma tissue [49]. Upon overexpression, it promoted cell proliferation of melanoma cells and regulated glycolysis and lactate production [49]. With an untargeted proteomics experiment, we identified, among others, *Pxk* as a potential sense target mRNA. PXK is a serine/threonine kinase involved in telomere length control. Its deregulation is a hallmark of cancer and it is influenced by c-Myc activity [50–52]. The ability of *circ5533* to increase PXK protein amount was then confirmed both on co-transfected cDNA and on the endogenous mRNA acting at the post-transcriptional level. To better understand the mechanism of action, we first excluded a sponge activity on endogenous miRNAs. We then analyzed its distribution in polysome profiling, finding that *circ5533* RNA is mainly associated with the 80S ribosome and polysomes fractions, proving it is acting at the level of translation. After showing the direct interaction between *circ5533* RNA and *Pxk* mRNA, we demonstrated that *circ5533*’s effect is the consequence of a SINEUP(IRES)-like mechanism by identifying BDs and TSs. The localization of TS1 at the 5’UTR of the target mRNA resembles the positions of TSs for natural and synthetic SINEUPs although it does not include the translation initiation codon AUG and its adjacent sequences. Furthermore, deletion analysis sustains a model where a simultaneous pairing could occur to both 5’ and 3’ UTRs of the target mRNA through two different BDs in *circ5533*. Taken together, these results suggest that the simultaneous pairing of *circ5533* at both 5’ and 3’ UTRs of *Pxk* mRNA could lead the target mRNA to assume a loop conformation to stimulate mRNA binding to the ribosome subunits, thus facilitating protein translation re-initiation. This hypothesis would be, indeed, consistent with what was observed in both cap-dependent or –independent translation initiation, where the ability of the poly(A) binding proteins to interact with cap or IRES-binding proteins can mediate circularization of the mRNA by linking the 5’ and 3’UTRs in a closed loop [53,54]. In addition, recent studies reported that circular RNAs can control gene expression by targeting and stabilizing specific mRNAs, thus regulating their translation [55]. This observation is likely consistent with our findings since the association of *circ5533* to polysomes could be a key factor for *Pxk* mRNA stabilization in the translation complex. However, how much IRES-like mechanisms, as defined in *cis*, are indeed used in *trans* is a fascinating topic of further investigation.

In summary, this study shows that natural SINEUPs include linear and circular transcripts with an embedded IRES sequence as ED. This highly heterogeneous class of regulatory RNAs may involve a variety of molecular mechanisms, mirroring the diversity found among IRES types in linear and circular protein-coding genes. Importantly, we show that SINEUP activity can be exerted on target mRNAs transcribed from *loci* located anywhere in the genome. Any experiment describing the transcriptome-wide repertoire of linear and circular non-coding RNA-mRNA heteroduplex in cells will increase our understanding of their pairing rules and better define the relevance of SINEUP(IRES)-like activities in cells’ homeostasis and disease.

## ACKNOWLEDGMENTS

We are indebted to all the members of the SG laboratory and of the RNA Initiative@IIT for thought-provoking discussions. We are grateful to the technical and administrative staff of IIT (especially Rosa Maria Cossu, Eva Ferri, Claudio Pisano, Elena Ghirelli, Fabrizio Torri and Alessandra Sanna), SISSA (Cristina Leonesi), and Università del Piemonte Orientale. We thank Prof Alberto Inga (University of Trento, Italy) for the plasmids used in bicistronic IRES assays, Prof. Marta Biagioli (University of Trento, Italy) for the pcDNA3.1(+) ZKSCAN1 MCS Exon circularizing vector, Prof Licio Collavin (University of Trieste, Italy) for HepG2 cells and Dr. Azzurra Codino for the help with DICER KO cells. We acknowledge that the research activity herein was carried out using the IIT HPC infrastructure.

## AUTHOR CONTRIBUTIONS

SD conceived, performed and analyzed the results of the experiments on *circ5533*, wrote the manuscript; ATM conceived, performed and analyzed the results of the experiments on IRES activity, *SINEUP(IRES)-DJ-1*; MV performed the bioinformatics analysis on circRNAs and RNA-seq experiments; BP carried out experiments on *circ5533* and wrote the manuscript; FA performed the bioinformatics analysis on circRNAs and RNA-seq experiments; PL carried out some bioinformatic analysis on circular RNAs; CaB carried out experiments and analyzed results; OP carried out experiments on *circ5533*; ClB carried out proteomics experiments and analyzed results; AA carried out proteomics experiments and analyzed results; MS carried out some experiments; DD carried out some preliminary experiments; VDC carried out experiments on *circ5533*; LB carried out some experiments and analyzed results; EB carried out some experiments and analyzed results; GGT analyzed results and discussed data; PC analyzed results and discussed data; CS conceived the project, analyzed the results and wrote the manuscript; FP conceived the project, analyzed the results and wrote the manuscript; LP carry out experiments on *dicer^-/-^* cells and analyzed results; SE conceived the project and analyzed results; SZ conceived and supervised the study; RS conceived and supervised the bioinformatics analysis, wrote the manuscript; SG conceived and supervised the study, wrote the manuscript.

## Material and Methods

### Plasmid Constructs

Synthetic SINEUP constructs were generated to assess the activity of both invSINEB2 and IRES elements in regulating target mRNA translations in *cis* or *trans*. SINEB2 elements (in direct and inverted orientation) were PCR-purified (primers are reported in Table S3), digested with EcoRI and XhoI restriction enzymes, and cloned into pRUF plasmid between Rluc and Fluc CDSs. Empty pRUF and pRUF-c-myc IRES, containing *c-myc* IRES between Rluc and Fluc sequences, were obtained from the laboratory of Prof Alberto Inga (University of Trento) and used as controls. Promoterless bicistronic vectors were generated by removing the SV40 promoter from the bicistronic vector pRUF and the pRUF vectors containing the putative IRES sequences by PCR amplification using divergent primers (Supplementary Table 3) and the CloneAmp™ HiFi PCR Premix (Takara) polymerase. To generate the constructs for the circular GFP reporter assay, the vector pcDNA3.1(+) ZKSCAN1 MCS-WT Split GFP + Sense IRES was purchased from Addgene (Plasmid #69909), linearized by PCR and using the divergent primers reported in Supplementary Table 3 and CloneAmp™ HiFi PCR Premix (Takara) polymerase. The primers were designed to linearize the plasmid and at the same time remove the sense EMCV IRES contained in the vector. The sequences containing the putative IRESs under investigation (*c-myc* IRES, invSINEB2) were amplified using as a template the respective pRUF plasmids with primers (Supplementary Table 3) containing an overhanging region (20 nt) overlapping with the linearized pcDNA3.1(+) ZKSCAN1 MCS-WT Split GFP at the desired site of insertion. SINEUP RNA-encoding plasmids were gene-synthesized using *Δ5’-ASUchl1* as the backbone sequence and cloned into pCS2^+^[2,3]. *Δ5’-AS Uchl1* (SINEUP-ΔBD) RNA lacks the overlapping region (BD) targeting *Uchl1* sense mRNA, and it comprises the inverted SINEB2 (ED) with a downstream Alu element and 3’ tail sequences. SINEUP-ΔBD was subcloned in pCS2^+^ from the pCDNA3.1(-) vector[3] using EcoRI and HindIII as restriction sites. SINEUP-ΔED, only retaining DJ-1 BD sequence, was PCR-cloned using primers reported in Table S3. *SINEUP-DJ-1* binding domain was designed, in antisense orientation, targeting endogenous *DJ-1* mRNA at position −40/+4 considering A, in the ATG start codon, at position +1[3]. miniSINEUPs, comprising a BD targeting mRNA of interest and an ED made by only the invSINEB2 sequence, were designed to target endogenous *DJ-1* or ectopic GFP mRNAs. *miniSINEUP-DJ-1* was PCR-cloned in pCS2^+^ vector using primers reported in Table S3. *miniSINEUP-GFP* was cloned into pCDNA3.1(-) plasmid as previously described^10^. To co-express exogenous GFP mRNA with miniSINEUP constructs, peGFP-C2 (Clonetech) was used. SINEUP (IRES) constructs are characterized by the presence of viral or cellular IRES elements as ED instead of canonical invSINEB2 sequence (Figure 3a-b); *DJ-1*-targeting BD was used. IRES sequences were gene-synthesized and subcloned in SINEUP-ΔED backbone between XhoI and XbaI restriction sites in the pCS2^+^ vector. Mutated invSINEB2 and HCV IRES sequences (Figures 3 and 4) were gene-synthesized and cloned into pCS2^+^-SINEUPΔED construct between XhoI and XbaI restriction sites. miniSINEUP (IRES)s exclusively comprises an IRES element as ED with different BDs targeting *DJ-1* or GFP mRNAs. *miniSINEUP (HCV IRES)-DJ-1*, *miniSINEUP (Polio IRES)-DJ-1* and *miniSINEUP (c-myc IRES)-DJ-1* were PCR-purified, using primers reported in Table S3, and cloned into pCS2^+^ at EcoRI and HindIII restriction sites. *miniSINEUP (c-myc IRES)-GFP* was generated by using XhoI and EcoRI restriction enzymes to replace DJ-1 BD with a GFP BD. To assess circSINEUP activity, *miniSINEUP-(c-myc IRES)-GFP* was sub-cloned in pcDNA3.1(+) ZKSCAN1 MCS Exon circularizing vector (kindly provided by Prof. Marta Biagioli, University of Trento, Italy) between ClaI and AgeI restriction sites, giving rise to pcDNA3.1(+) ZKSCAN1-circSINEUP(*c-myc* IRES)-GFP. Studies on the role of *circ5533* working as potential circSINEUPs were carried out by cloning the reported *circ5533* sequence (<u>hsa_circ_0085533</u> Circular RNA Interactome) in pCDNA3.1(+)LACCASE2 MCS Exon Vector (#69893, Addgene) or pCDNA3.3(+)ZKSCAN1 MCS Exon Vector (#69901, Addgene); both vectors can express circular RNAs of specific sequences in mammalian cells. Constructs were purchased from GeneScript, the MCS was completely removed in pcDNA3.1(+)LACCASE2 MCS Exon Vector – *circ5533* to properly circularize the insert. Empty pcDNA3.1(+)LACCASE2 MCS Exon Vector and pcDNA3.1(+)ZKSCAN1 MCS Exon Vector were used as controls.

*circ5533* ΔBD1 (deleted BD1) or *circ5533* ΔBD2 (deleted BD2) in pcDNA3.1(+)LACCASE2 MCS Exon Vector were gene-synthesized. WT *PXK* mRNA (NM_017771.5, NCBI) was tagged with a FLAG sequence and cloned into pcDNA3.1(+) vector at the NheI and ApaI restriction enzyme sites. In addition, putative target sites (TS1, TS2, TS3 at the 5’UTR; TS4 and TS5 at the 3’UTR) were individually deleted from the respective UTRs in *Pxk* mRNA. PXK-FLAG ΔTS1 (deleted TS1), PXK-FLAG ΔTS2 (deleted TS2), PXK-FLAG ΔTS3 (deleted TS3), PXK-FLAG ΔTS4 (deleted TS4) and PXK-FLAG ΔTS5 (deleted TS5) were respectively cloned in pcDNA3.1(+) vector. Potential *Pxk* target sites at the 5’UTR (TS1, TS2, TS3) were individually cloned immediately upstream PXK-FLAG sequence (without 5’ and 3’UTRs) in pcDNA3.1(+) plasmid, while target sites at the 3’UTR (TS4 and TS5) were added downstream of PXK-FLAG CDS after the STOP codon. All constructs were procured from GeneScript.

### Cell Lines and Transfection Conditions

HEK 293T/17 and U2OS cells were obtained from ATCC (Cat. No. ATCC-CRL-11268 164 293T/17; Cat. No. HTB-96). Both cell lines were maintained in culture with Dulbecco’s Modified Eagle Medium (GIBCO) supplemented with 10% FBS (SIGMA) and 1% antibiotics (penicillin/streptomycin), as suggested by the vendor. HepG2 cells were kindly provided by Professor Collavin L. from the University of Trieste (Italy)[56] and maintained in culture with Eagle’s minimal essential medium (SIGMA) supplemented with 10% FBS, 1% antibiotics, 1% GlutaMAX and 1% non-essential amino acids. Dicer KO mESC cells were a kind gift from Gregory Hannon (CRUK Cambridge Institute, UK), and were maintained in 15% fetal calf serum, 2 mM glutamine, 1 mM sodium pyruvate, 1 mM non-essential amino acids, 0.05 mM β-mercaptoethanol, 100 U/mL penicillin/streptomycin, and 1000 U/mL recombinant mouse LIF (Invitrogen).

The experiments in all cell types were performed with Fugene HD (Roche) transfection reagent, following the manufacturer’s instruction. In detail, cells were seeded in 6-well plates the day before transfection at 60% confluence and transfected with SINEUP or SINEUP(IRES) or dual-luciferase plasmids using Fugene HD reagent in the ratio of 1:3 (w/v) with a total of 2µg plasmid DNA transfected per well. Bicistronic mRNA was transfected in HEK293T cells with Lipofectamine^TM^ MessengerMAX^TM^ Transfection Reagent (Thermo Fisher Scientific, LMRNA015). For circular GFP experiments, pCDNA3.1(+)ZKSCAN1-split GFP vectors were transfected in HEK293T cells using Lipofectamine 2000 (Invitrogen) as a transfecting reagent. For the SINEUP-activity assay, experiments were performed as previously described[3]. Cells were harvested after transfection in two equal parts for western blot and qPCR analysis for each replicate. To evaluate circSINEUP-GFP activity, HEK293T cells were seeded in 6-well plates and transfected with a total of 1 µg of plasmid DNA per well using Lipofectamine 2000 (Invitrogen) as transfecting reagent. We co-transfected the constructs of interest with pEGFP, in a 1:6 ratio (pEGFP:SINEUP/circSINEUP constructs) as previously described[3]. Cells were harvested at 48 h after transfection and divided into two equal parts for fluorescence assay and qPCR analysis for each replicate.

For *circ5533* experiments, HEK293T cells were seeded in 6-well plates and transfected with 1µg of plasmid DNA per well using Lipofectamine 2000 (Invitrogen) as transfecting reagent. Co-transfections were carried out using constructs of interest in a 1:12 ratio (PXK-constructs:*circ5533*). Cells were collected after 48 h in two equal parts for western blot and qPCR analysis for each replicate.

### Dual Luciferase Assays in Cell Lines (plasmids and IVT mRNAs)

Empty and *c-myc*-IRES pRUF plasmids were kindly provided by Professor Alberto Inga from the University of Trento (Italy). Cells were plated the day before transfection as previously described. HEK293T cells were transfected with empty, invSINEB2-, dirSINEB2-, and c-myc-IRES-containing pRUF plasmids. Cells were harvested at 48 h after transfection. For dual Luciferase assay with bicistronic mRNA, pRUF vectors of interest were linearized with XbaI enzyme. *In vitro* transcription of bicistronic mRNA was performed by using mMESSAGE mMACHINE^TM^ T7 ULTRA Transcription Kit (Thermo Fisher Scientific, AM1345). Six hours following transfection, cells were harvested. Both DNA and RNA luminescence assays were performed using the Dual-Glo Luciferase assay system (Promega) following the manufacturer’s instructions. Luminescence was measured using the microplate reader INFINITE 200 PRO (TECAN). The empty pRUF (with NO IRES between the two CDS) luminescence level was used to normalize the transfection efficiency or background effects as previously described^39,40^. IRES activity was calculated as the ratio of firefly luciferase (Fluc) activity to renilla luciferase (Rluc) activity. IRES activity control experiments were performed to exclude any cryptic promoter or alternative splicing events. RNA was purified from harvested cells and used for RT-PCR. All experiments were performed in duplicate for at least four independent biological replicates.

### Translatable circular mRNA assay

HEK293T cells were plated the day before transfection. Non-transfected cells and HEK293T cells over-expressing pCDNA3.1(+)ZKSCAN1 MCS-WT Split GFP + Antisense IRES (#699910) were used as negative controls. pCDNA3.1(+)ZKSCAN1 MCS-WT Split GFP + *cmyc* IRES (see “plasmid constructs”) was used as positive control. The potential IRES-like activity of invSINEB2 element was investigated by overexpressing pCDNA3.1(+)ZKSCAN1 MCS-WT Split GFP + invSINEB2 (see “plasmid constructs”) in HEK293T cells. Cells were harvested 48h following transfection. The Percentage of GFP-positive cells was calculated through Sony SH800 CellSorter.

### RNA Isolation, Reverse Transcription and Quantitative Real Time-PCR (qRT-PCR)

Total RNA was extracted with RNeasy Mini kit (Qiagen, #74106) according to the manufacturer’s protocol. On-column DNAse digestion was performed for 30 min at room temperature. 1 μg of total RNA was reverse-transcribed using iSCRIPT cDNA Synthesis Kit (Bio-Rad) according to the manufacturer’s protocol. Each qPCR Real-time reaction was carried out in a total volume of 10 µL with 20 ng of cDNA using SYBR-Green PCR Master Mix (BioRad) and an iCycler IQ Real-time PCR System (Bio-Rad). qPCRs were performed in the following conditions: 3 min at 95 °C and 40 cycles of 10 s at 95 °C plus 5 s at 60 °C. The specificity of PCR amplicons was checked with melting curve analysis and RT-samples. Values were normalized to *GAPDH* mRNA levels, the fold change was calculated using the 2^-ΔΔCt^ method.

For circRNA extraction, RNA was purified using miRNeasy Mini Kit (Qiagen, #74106). For PXK-FLAG experiments qPCR analysis was performed with two different primer pairs: one to detect endogenous PXK mRNA and one to specifically detect PXK-FLAG mRNA.

All primers used in this study are reported in Table S4.

### Western Blot Analysis

Harvested cells were lysed in radioimmunoprecipitation assay (RIPA) buffer with the addition of protease inhibitor cocktail (Sigma-Aldrich, Cat. No. P83490), briefly sonicated, and boiled with 1X Laemmli Buffer for 5 min at 95 °C. Then, 5 μg of total lysate, was resolved by 10% or 4-20% SDS-PAGE TGX precast gels (Biorad) and transferred to nitrocellulose membrane using Trans-Blot Turbo Transfer System (Bio-Rad). Membranes were blocked with 5% non-fat dry milk in TBS/0.1% Tween 20 and incubated with appropriate primary antibody. All antibodies used in this study are listed in Table S5. Proteins of interest were visualized with SuperSignal West Pico PLUS Chemiluminescent Substrate (Thermo Fisher Scientific, Cat. No. 34579). Western blot images were acquired with ChemiDoc MP Imaging System (Bio-Rad) and band intensity was calculated using ImageJ Software.

### RNA Pull-Down (RPD)

The PFA-crosslinked RNA pull-down protocol was performed as described in [57] with some modifications according to Bevilacqua *et al.* [58]. HEK293T cells were seeded in 100 mm dishes and transfected with 16µg of plasmid DNA per dish using Lipofectamine 2000 (Invitrogen) as transfectant. After 48h, cells were fixed with 1% PFA for 10’ at room temperature. PFA was quenched adding 1/10 volume of glycine 1.25 M for 5’ at RT. Following the treatment, cells were collected and lysed. The lysate was incubated at 95°C for 3’, moved on ice to cool down and finally divided into two aliquots per sample. For each sample, one aliquot was added to 500 pmol of mixed probes while the other was added to water. Samples were then incubated at 65°C for 5’and then added an equal volume of pre-heated (65°C) Binding Buffer (10mM Tris-HCl 7.5, 1mM EDTA, 2M NaCl). From each sample, 1/10 of the total volume was utilized as input. Samples and probes were incubated for 4h at RT on a tube rotor. Finally, Dynabeads^TM^ MyOne^TM^ Streptavidin C1 (Thermo-Fisher Scientific, 65001) were added following the manufacture’s protocol. Elution was carried out by adding TRIZOL reagent and RNA extraction performed as previously described for total RNA. RNA was retro-transcribed and analyzed by qRT-PCR.

### Polysome profile analysis

Polysome profiling was carried out according to protocol reported in [59]. Cells were cultured and transfected in 150 mm dishes. Before harvesting, cells were incubated with cycloheximide (Sigma-Aldrich, 01810), at a final concentration of 100 µg/ml, for 10 minutes at 37 °C. Then, cells were washed 3 times with cold PBS supplemented with 100 µg/ml CHX, collected and lysed in 400 µL of polysome extraction buffer (PEB)[59] containing 100 µg/ml CHX, 1:1000 RiboLock RNase inhibitor (Thermo-Fisher Scientific, EO0382) and 1X protease inhibitors (Sigma-Aldrich, P8340). Lysates were loaded on 15-50% sucrose density gradients and centrifuged (Beckman ultracentrifuge) for 1 hour and 30 minutes in a SW41Ti swinging bucket rotor at 39000 rpm at 4 °C. Gradient fractions were recovered in 28 fractions using Piston Gradient Fractionator (Biocomp Instruments) and UV absorbance was continuously measured (260 nm) using Triax flowcell. RNA from each fraction was first precipitated in EtOH/C_2_H_3_NaO_2_ and then purified through RNA Clean & Concentrator-5 kit following the manufacturer’s protocol (Zymo Reaserach, R1016). RNA distribution profiles were analyzed by qRT-PCR.

### Fluorescence-based Assay

First, 100 µL of harvested cells suspension in PBS were seeded in a 96-well microplates; then, 100 µL of PBS-diluted NucBlue Live Cell Stain Reagent (Invitrogen, R37605) was added. GFP fluorescence was measured with Spark Microplate Fluorescence Reader (TECAN Systems) setting up the excitation wavelength at 485 nm and the emission wavelength at 535 nm. GFP fluorescence intensity was normalized on NucBlue fluorescence, measured with an excitation wavelength of 360 nm and an emission wavelength of 465 nm.

### Statistical Analysis

All data are expressed as mean ± standard deviation on n ≥ 3 replicates. Statistical analysis was performed with GraphPad Prism software. Statistically significant differences were assessed with Student’s t-test, one-sample t-test or one-way ANOVA test.

### Bioinformatics analysis for circRNA identification

For circRNAs analysis, we used the circBase database[60]. Nucleotide sequences of circRNAs are available for human, mouse, *Caenorhabditis elegans*, *Latimeria chalumnae*, and *Latimeria menadoensis*. We downloaded the human circRNA sequences from http://www.circbase.org/download/human_hg19_circRNAs_putative_spliced_sequence.fa.gz. IRES annotation and sequences were downloaded as supplementary material from [42]. The same group also developed a machine learning approach to predict IRES activity and explore the relationship between RNA sequences and IRES activity[43]; a complete table of the 55,000 oligos tested by the group, together with the scores calculated for the promoter activity and splicing were downloaded from https://bitbucket.org/alexeyg-com/. Starting from this table, entries belonging to Human_5UTR_Screen, IRESite_blocks and High_Priority_Genes_Blocks were considered. In addition, other selected filters were as follows: promoter_activity < 0.2, splicing_score > −2.5, ires_activity > 206.3 (based on eGFP expression) as discussed in [42]. A total of 570 entries passed these filters: 254 entries belong to the Human_5UTR_Screen subset, 292 to High_Priority_Genes_Blocks and 24 to IRESite_blocks. The obtained IRES sequences were trimmed at both ends and then aligned against circRNA sequences from CircBase using BLAST[61] with the following parameters: word size 11, e-value 100 with selected soft masked. Entries with 100% IRES coverage (*qcovhsp*), 0 gaps and a maximum of 1 mismatch were considered.

The Virtual Ribosome tool (Dna2pep v1.1)[44] was used to predict the longest complete ORF for each circRNA by searching across all positive reading frames with methionine as start codon and a canonical stop (parameters: -o strict, -r plus).

### Mass Spectrometry

2 × 10^5^ HEK293T cells were seeded in 6-well plates and transfected with 1µg of pcDNA3.1(+) ZKSCAN1 MCS Exon *circ5533* to over-express *circ5533* RNA. Cells transfected with empty pcDNA3.1(+) ZKSCAN1 MCS Exon were used as control. Cells were harvested at 48 h after transfection. RNA and proteins were extracted from the same sample for each replica. A total of 6 replicates per condition were prepared. Protein fractions were then analyzed by performing mass-spectrometry-based untargeted proteomics experiments. Protein concentration for each sample was assessed using a standard protein assay kit (Invitrogen, Cat. 23225).

The volume corresponding to 50 µg of proteins was then collected from each sample to be digested before analysis with mass-spectrometry coupled to liquid chromatography (LC-MS/MS) analysis. The disulfide bonds on cysteine residues were reduced with 10 µL of 100 mM dithiothreitol (DTT) at 56 °C for 30 min; cysteine residues were then alkylated with 30 µL of 100 mM iodoacetamide (IAA) for 20 min in the dark. The reducing and alkylating solutions were then removed by precipitating the protein content in cold acetone overnight at −20 °C. The following day, the samples were centrifuged (20000xg for 30 min at 4 °C), the supernatant was discarded, and the protein pellets were dried under a nitrogen stream. Dried pellets were then re-dissolved in 100 µL of digestion buffer (50 mM NH_4_HCO_3_ in MilliQ water, pH=8) before adding 1 µg of trypsin for overnight digestion at 37 °C. The resulting peptides were dried under vacuum and then dissolved in 150 µL of 3% Acetonitrile (ACN) + 0.1% Formic Acid (FA) for LC-MS/MS analysis.

### LC-MS/MS and data analysis

The same amount of tryptic peptides from each sample (1.66 µg), was injected on a NanoAcquity chromatographic system and analyzed with a TripleTof 5600+ mass spectrometer equipped with a NanoSpray III ion source. The trapping phase was performed on a 180 µm x 20 mm Acquity C18 column for 4 min at 4.0 µL/min flow rate in 1% ACN + 0.1% FA. The peptides were then moved and separated using a PicoFrit C18 column (75 μm x 25 cm, from NewObjective Inc., Woburn, MA, USA); the elution of the peptides was performed at 300 nL/min with a 2 h gradient of ACN in water (3% to 45%, both added with 0.1% FA). ACN content in the eluents was then increased to 90% in 5 min and kept at 90% for an additional 5 min to wash the column. The system was then re-equilibrated to 3% ACN for 18 min. The peptides were analyzed in positive ion mode with the following parameters: ion spray voltage at 3050 V, spray gas 1 kept at 16, curtain gas at 30, and declustering potential at 80 V. Source temperature was set at 90 °C. For protein quantification, the mass spectrometer was operated in data-independent acquisition (DIA) mode, following the SWATH protocol for label-free proteomics[62]. For SWATH analysis, precursor ions were selected within the 400-1250 m/z range, with a variable window width from 7 to 50 Da. After a full range survey scan of 250 ms, 100 consecutive SWATH experiments (100-1500 m/z) were performed, each lasting 25 ms. For protein quantification, the spectra were searched against a modified version of the PanHuman ion library[63] using only no-shared peptides. For the quantification, the following settings were used: minimum peptide confidence of 90%, 50 ppm maximum mass tolerance, 30 min maximum RT tolerance, 6 MRM transitions per peptide, and modified peptides were not allowed. With these settings, a total of 4921 proteins were quantified in the samples. For statistical analysis, raw data were imported in MarkerView software and normalized using the Most Likely Ratio (MLR) method[64]. To select only the significantly (p.value < 0,05) dysregulated proteins three un-paired, two-tailed t-tests were performed (comparing cir5533 vs Empty).

### RNA-seq analysis

pcDNA3.1(+) ZKSCAN1 MCS Exon *circ5533* and relative control were transfected as described above. Total RNA and proteins were extracted from the same sample for each replica. RNA fractions were analyzed with RNA-seq. The RNA-seq libraries were prepared starting from 200 ng of DNase-treated total RNA according to the Illumina TruSeq Stranded Total RNA protocol (Document # 1000000040499 v00, © 2021 Illumina, Inc.). The libraries qualities were assessed by running a 2100 Bioanalyzer DNA 1000 chip (Agilent Technologies, Inc). A 1.4 nM pool of each QC passed RNA-seq library was sequenced by running a Single Read (SR) 100bp NovaSeq 6000 S2 flow cell on the NovaSeq 6000 Sequencing System (Illumina, Inc.). The fastq files were obtained by demultiplexing the bcl basecall files using the 1.8.4 version bcl2fastq script (Illumina, Inc.). Reads mapping was performed against the reference genome (GRCh38, Ensembl release 98) using STAR[65] (v. 2.6) . The tool *featureCounts*^71,74^ from the Subread package (v. 2.0.0) was used to count the number of reads for each gene. Finally, EdgeR[67] (v. 3.26.8) was used to perform the DEG analysis with default parameters using an FDR cutoff of 0.05.

### Bioinformatics analysis of BD sequences

UniProt accessions of upregulated proteins (resulting from the proteomics) were mapped to Ensembl transcript IDs through the UniProt ID mapping tool. Due to the one-to-many relationship of this data type given by the different isoforms that each gene can produce, this mapping resulted in a total of 304 transcripts (one UniProt ID to many Ensembl transcript IDs). The transcripts were aligned against *circ5533* RNA sequence using blastn in BLAST[61] (v2.7.1, parameters: -word_size 5; -evalue 999999; -dust no; -soft_masking false; -outfmt “6 qaccver saccver pident length mismatch gapopen qstart qend sstart send evalue bitscore frames qseq sseq btop”, -num_threads 12). The resulting table was parsed on the frames column to keep matches of the 1/-1 type (i.e. in opposite orientation). In addition, we parsed the btop column defined as “Blast trace-back operations” (which describes the entire alignment produced by BLAST in one string) to filter out results presenting perfect matches shorter than 8 nts. In order to annotate the position of each resulting match among the different isoforms, genomic UTRs and CDS coordinates were converted to local coordinates with a custom-made script in R[68] (v. 3.5.1) using the Ensembl human gene set annotations (release 98). RepeatMasker (v.4.0.7, RepBase Update v20181026) was run with default parameters to search for repeats in the *circ5533* sequence. This search resulted in the annotation of two simple_repeats (pos: 182-243; 667-694 of *circ5533)*. Finally, the potential BD coordinates resulted from the analysis described above were compared with these repeat coordinates in order to exclude any overlap.

## SUPPLEMENTAL FIGURES

**Figure S1. Related to Figure 1.**
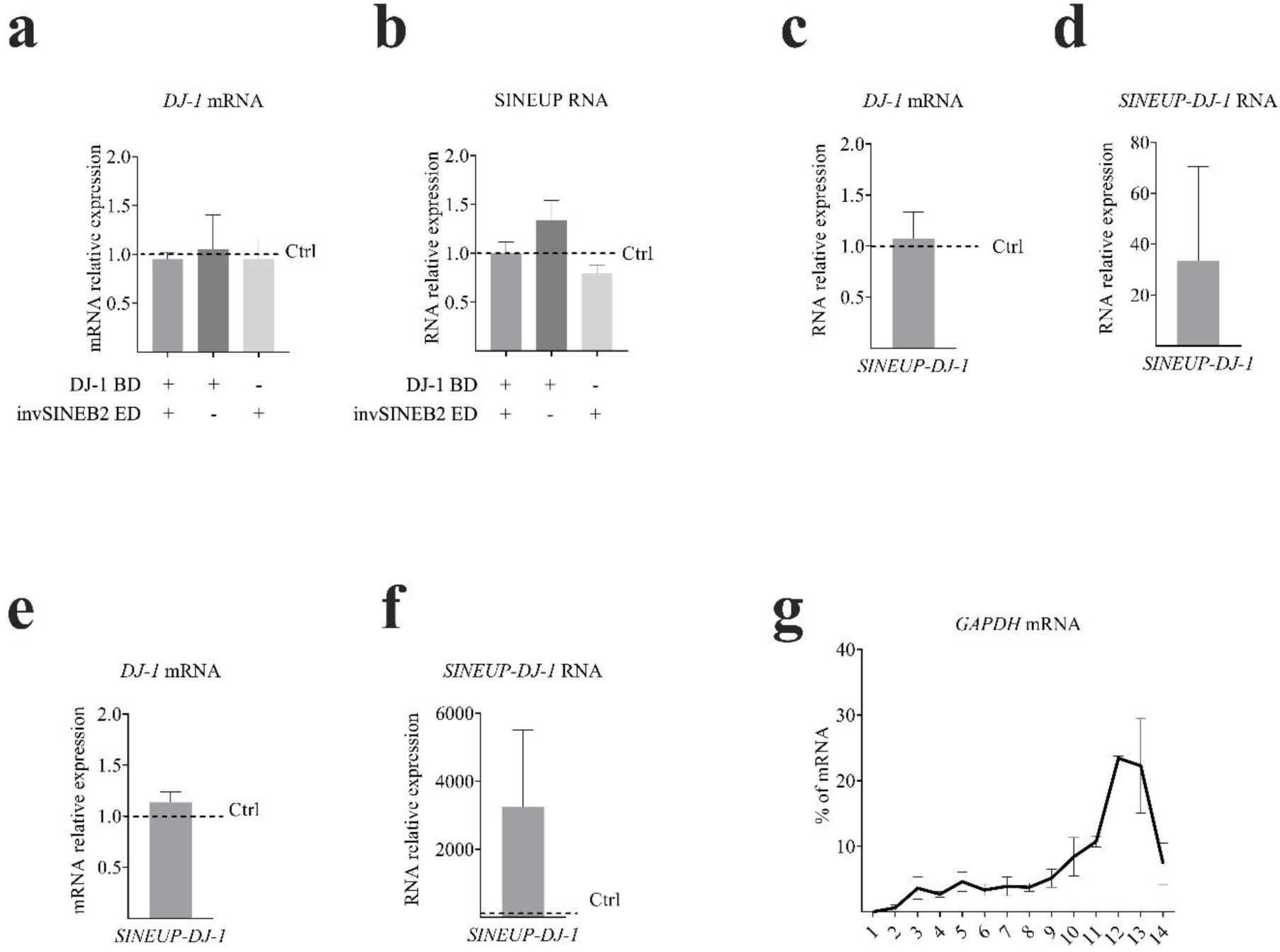
**(a-b) *SINEUP DJ-1* activity.** RNA levels. Canonical *SINEUP-DJ-1* was tested along with mutated forms. Total RNAs were extracted at 48 h after HEK293T cell transfection, reverse-transcribed and analyzed through real time PCR. **a)** DJ-1 mRNA relative expression was quantified and reported on the graph using empty pCS2+ as negative control (Ctrl=1, dotted line). *GAPDH* mRNA was used to normalize. No significant variation was observed (ordinary one-way ANOVA, Holm-Sidak’s multiple comparisons test) **b)** Expression of mutated constructs was also assessed through Real-time PCR. SINEUP mutant RNA expression was quantified compared to *SINEUP-DJ-1* RNA levels. All plots indicate mean ± SD, and representation of n = 3 independent biological replicates. **(c-e) SINEUP RNA pull-down. c)** *DJ-1* mRNA expression for RPD. **d)** *SINEP-DJ-1* expression for RPD. **e)** *DJ-1* mRNA expression for ribosome fractionation. **(f-g) Ribosome fractionation. f)** *SINEP-DJ-1* expression for ribosome fractionation. DJ-1 mRNA and *SINEP-DJ-1* RNA relative expression was quantified and reported on the graph using empty pCS2+ as negative control (Ctrl=1, dotted line). *GAPDH* mRNA was used to normalize. **g)** Lysates from control and treated cells were fractionated by sucrose gradients and separated through a UV detector continuously measuring the solution absorbance (260 nm). The relative distributions (in %) of *GAPDH* mRNA were analyzed by RT-qPCR in each of the 13 gradient fractions. RNA distribution was reported as the mean ± SD of 3 replicates.

**Figure S2. Related to Figure 2.**
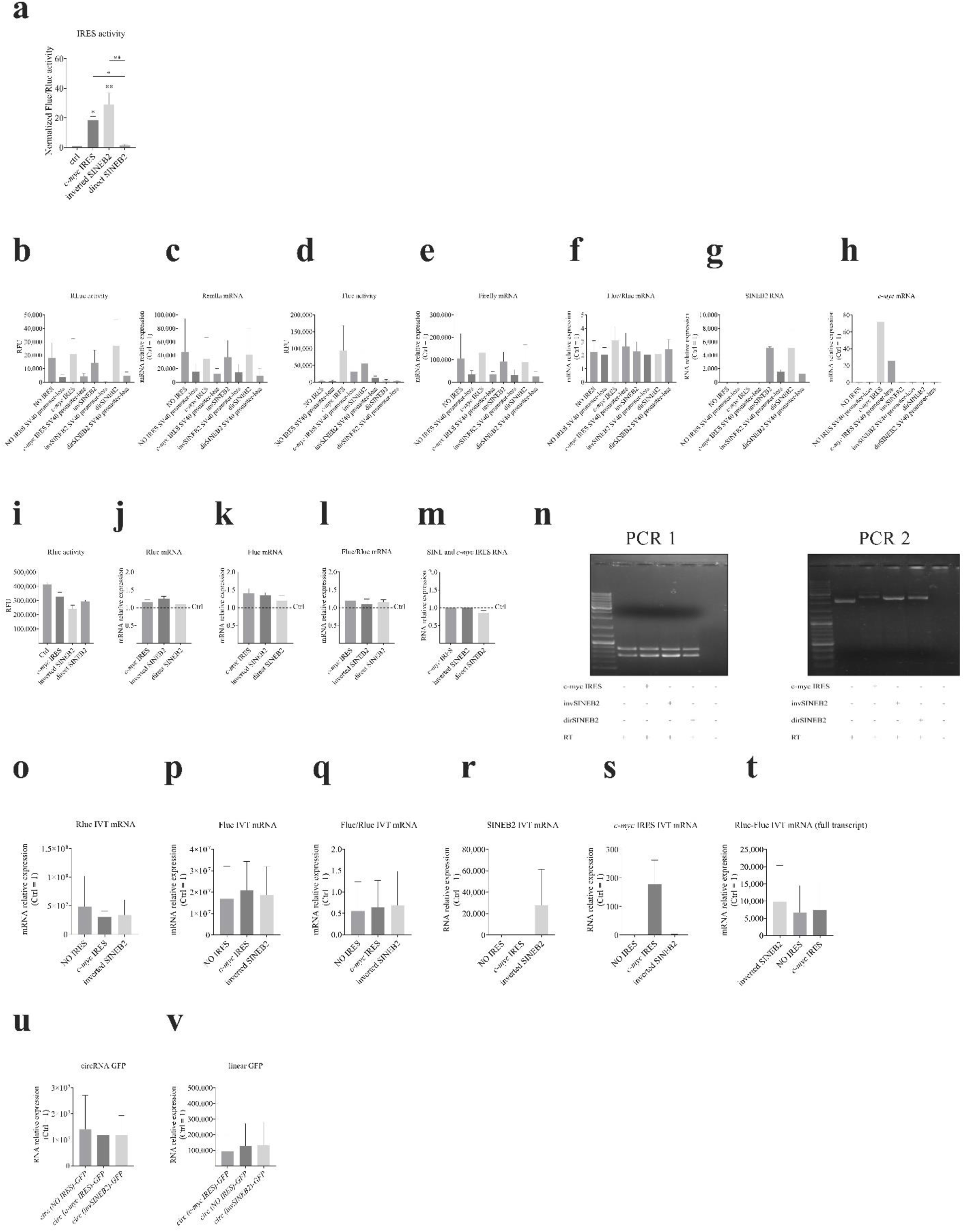
**a) IRES activity assay in U2OS cells.** IRES activity. SINEB2 element, in direct (dirSINEB2) and in inverted (invSINEB2) orientations, and *c-myc* IRES were separately cloned between Rluc and Fluc sequences in pRUF-dual luciferase reporter vectors. Empty pRUF-MCS and pRUF-*c-myc* IRES were respectively used as negative and positive controls. Luciferase intensities were measured at 48 h after U2OS cell transfection of each pRUF dual luciferase reporter plasmid (reported in Figure1). Results are shown as the detected intensity ratio of Fluc and Rluc (Fluc/Rluc) normalized against the empty pRUF-MCS background signal. IRES activity was calculated compared to the negative control (Ctrl=1, dotted line). Reported graph shows that invSINEB2 induced Fluc protein translation *in cis* at higher levels than the positive control. Plot indicates mean ± SD of 4 biological replicates performed in duplicate; *p <0.05, **p <0.01 were considered significant. Ordinary one-way ANOVA with Holm-Sidak’s multiple comparisons test was used to calculate significances. **(b-h) IRES activity (promoter-less plasmids).** Empty pRUF-MCS and pRUF-*c-myc* IRES, both SV40 promoter-less were used as negative controls. **b)** Rluc expression was quantified and showed on the graph as mean luminescence intensity (RFU) of n = 3 independent replicates. **c)** Rluc mRNA relative expressions were quantified and reported on the graph using non-transfected cells (only LIPO) as control = 1. **d)** Fluc activity was plotted on the graph as mean luminescence intensity (RFU) of n = 3 independent replicates. Expression of Fluc protein was calculated compared to the LIPO control. **e)** Rluc mRNA levels were quantified and reported on the graph using, as control, non-transfected cells. **f)** Fluc mRNA levels relative to Rluc expression (Fluc/Rluc) were reported on the graph. **g and h)** qRT-PCR assessed the expression of the inserted invSINEB2, dirSINEB2 and *c-myc*-IRES sequences. RNA levels were normalized on *GAPDH* mRNA expression. All plots indicate mean ± SD, and represent n=4 independent replicates. **(i-n) IRES activity (bicistronic plasmids).** SINEB2 element, in direct (dirSINEB2) and in inverted (invSINEB2) orientations, and *c-myc* IRES were separately cloned between Rluc and Fluc sequences in pRUF-dual luciferase reporter vectors. Empty pRUF-MCS and pRUF-*c-myc* IRES were used as negative and positive controls respectively. **i)** Rluc activity was quantified and reported on the graph as mean luminescence intensity (RFU) of n = 3 independent replicates. **(j-k)** Total RNAs were extracted at 48 h after HEK293T cell transfection, reverse-transcribed, and analyzed with real-time PCR. **j)** Rluc mRNA relative expressions were quantified and reported on the graph using empty pRUF as control (Ctrl=1, dotted line). **k)** Relative expressions of Fluc mRNAs were calculated compared to the empty control. **l)** Fluc mRNA levels relative to Rluc expression (Fluc/Rluc) were reported on the graph assessing the absence of cryptic promoters and thus of shorter transcripts containing only Fluc or Rluc sequences. No significant variation was observed. **m)** qRT-PCR confirmed the expression of the inserted *c-myc*-IRES, invSINEB2 and dirSINEB2 sequences. RNA levels were normalized on *GAPDH* mRNA expression. All plots indicate mean ± SD, and represent n=4 independent replicates. **n)** Specific region of cDNAs, derived from extracted RNAs, were amplified by performing two different PCRs (PCR1 and PCR2). For PCR1, a pSV40 forward primer (annealing downstream the TSS and upstream Rluc CDS) was used in combination with a reverse primer targeting Rluc CDS. PCR1-derived amplicons were loaded on an agarose gel and images captured. Double bands represent long and short amplicons made from unprocessed and processed transcripts respectively as a consequence of the presence of an intronic sequence downstream of the pSV40 promoter of the pRUF vector backbone. PCR2 was performed using the same pSV40 forward primer used in PCR1 in combination with a reverse primer annealing to Fluc CDS. Derived amplicons were visualized on agarose gel. All derived amplicons showed the expected size confirming the transcription of a unique “bicistronic” RNA for each construct. **(o-t) IRES activity (IVT mRNA).** SINEB2 element, in inverted (invSINEB2) orientation, and *c-myc* IRES were separately cloned between Rluc and Fluc sequences in pRUF-dual luciferase reporter vectors. Vectors were linearized and derived mRNAs *in vitro* transcribed. Bicistronic mRNA with no IRES sequences and with *c-myc* IRES were used as negative and positive controls respectively. Total RNA was extracted at 6 h following HEK293T cell transfection, reverse-transcribed, and analyzed with real-time PCR. **o)** Rluc mRNA relative expression was reported on the graph using non-transfected cells as control (Ctrl=1). **p)** Relative expression of Fluc mRNA was analyzed as previously described for Rluc mRNA. **q)** Fluc mRNA levels relative to Rluc expression (Fluc/Rluc) were reported on the graph to assess the same mRNA expression levels for the two CDSs. **r and s)** qRT-PCR confirmed the expression of the inserted invSINEB2 and *c-myc*-IRES elements. RNA levels were normalized on *GAPDH* mRNA expression. All plots indicate mean ± SD, and represent n=4 independent replicates. **t)** Rluc-Fluc whole bicistronic mRNA levels were reported on the graph. *GAPDH* was used to normalize.**(u-v) circRNA expression (circularizing vector).** GFP-circularizing vectors were transfected in HEK293T cells. 48 h after transfection, cells were harvested and RNA extracted. Non-transfected cells were used as control (Ctrl=1). Circular GFP with IRES in antisense orientation was the negative control while circGFP with an embedded *c-myc* IRES was the positive control. RNAs were extracted, reverse transcribed and analyzed with real time PCR. *GAPDH* was used to normalize. **u)** Expression of circular GFP was confirmed in all constructs. **v)** Linear *GFP* mRNA relative expressions were quantified and reported on the graph. As expected, linear counterpart resulted much lower compared to the circRNA over-expression. All graphs indicate mean ± SD, and represent n=3 independent replicates.

**Figure S3. Related to Figure 3.**
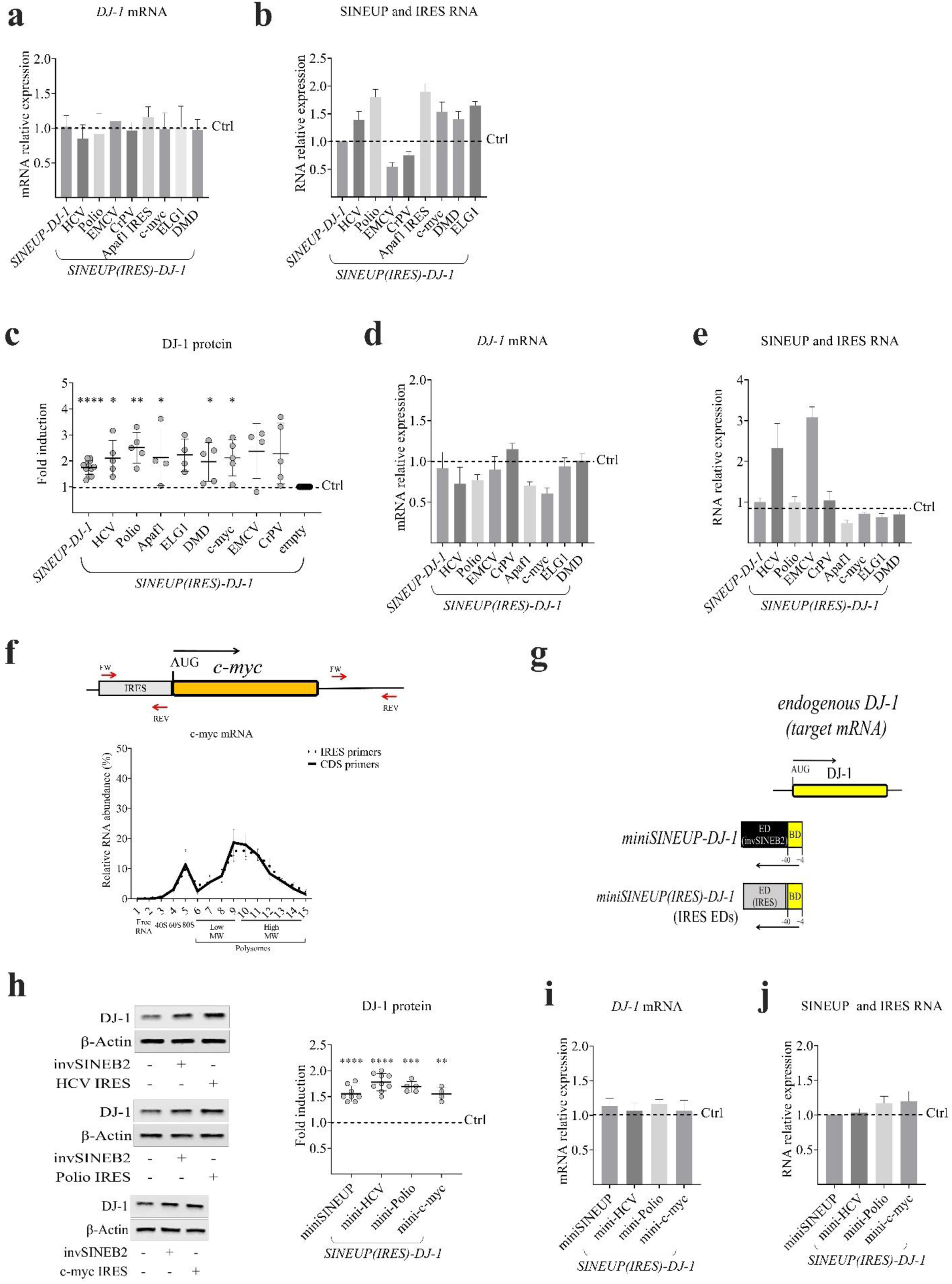
**a-b) *DJ-1* mRNA and SINEUP RNA levels.** *SINEUP(IRES)-DJ-1* comprises a BD, targeting *DJ-1* mRNA (−40/+4) in an antisense orientation, and different IRES sequences (viral or cellular) acting as potential EDs. SINEUP(IRES) constructs were transiently expressed in HEK 293T cells. Total RNAs were extracted at 48 h after transfection, reverse transcribed and analyzed with real time PCR. RNA levels were normalized on *GAPDH* mRNA. Empty pCS2+ was used as control (Ctrl=1, dotted line). **a)** *DJ-1* mRNA relative expressions were quantified and reported on the graph. No significant variations were observed. **b)** qRT-PCR confirmed the expression of all transfected *SINEUP(IRES)-DJ-1* constructs. All plots indicate mean ± SD, and represent n=4 independent replicas. (**c-e) *SINEUP(IRES)* activity in HepG2 cells. c)** Reassuming plot reporting *SINEUP(IRES)-DJ-1* activity. SINEUP constructs were transiently expressed in HepG2 cells. SINEUP activity was measured 48 h after transfection with western blot. Western blot analysis was carried out using anti-DJ-1 Ab to detect and quantify DJ-1 proteins. β-actin was used to normalize. Empty pCS2+ was used as control (Ctrl=1, dotted line). All SINEUP(IRES) RNAs increased DJ-1 protein levels in a canonical SINEUP-like mechanism. Plot reports DJ-1 fold induction mean ± SD; it represents at least 4 independent biological replicates; *p <0.05, **p <0.01, and ****p <0.0001 were considered significant (ordinary one-way ANOVA, Holm-Sidak’s multiple comparisons test). **d)** RNA levels. Total RNAs were extracted, reverse-transcribed, and analyzed with real time PCR. RNA levels were normalized on *GAPDH* mRNA. Empty pCS2+ was used as control (Ctrl=1, dotted line). *DJ-1* mRNA relative expressions were quantified and reported on the graph. No significant variations were observed. **e)** Expression of all transfected *SINEUP(IRES)-DJ-1* constructs were also assessed with real time PCR. All plots indicate mean ± SD and represent n=4 independent replicates. **f) Endogenous *c-myc* mRNA polysome distribution.** The relative distribution (in %) of *c-myc* mRNA, in non-transfected cells, was analyzed through Real-Time PCR using two different oligo pairs: one pairing at the level of IRES element and one at the 3’UTR of *c-myc* mRNA. RNA distributions were reported as the mean ± SD of 3 replicates. **(g-j) *miniSINEUP(IRES)* activity in HEK293T cells. g)** Schematic overview of *miniSINEUP-DJ-1* and *miniSINEUP(IRES)-DJ-1*. Mini synthetic SINEUP targeting *DJ-1* mRNA (yellow) comprising exclusively a BD (yellow) and an invSINEB2 as ED (black). *miniSINEUP(IRES)-DJ-1* comprises a BD, targeting *DJ-1* mRNA (−40/+4) in an antisense orientation, and different IRES sequences (viral or cellular) acting as potential EDs (light grey). **h)** Representative western blot images and reassuming plot reporting *miniSINEUP(IRES)-DJ-1 activity*. miniSINEUP constructs were transiently expressed in HEK 293T cells and their activity measured 48 h after transfection by western blot. Western blot analysis was carried out using anti-DJ-1 Ab to detect and quantify DJ-1 proteins. β-actin was used to normalize. Empty pCS2+ was used as control (Ctrl=1, dotted line). As expected, all miniSINEUP(IRES) RNAs increased DJ-1 protein levels. Plot reports DJ-1 fold induction mean ± SD; it represents at least 4 independent biological replicates performed in duplicate; ****p <0.0001 was analyzed with ordinary one-way ANOVA, Holm-Sidak’s multiple comparisons test. **i)** RNA levels. Total RNAs were extracted, reverse transcribed and analyzed with real time PCR. RNA levels were normalized on *GAPDH* mRNA. Empty pCS2+ was used as control (Ctrl=1, dotted line). *DJ-1* mRNA relative expressions were quantified and reported on the graph. No significant variations were observed. **j)** Expression of all transfected miniSINEUP(IRES) constructs was assessed with real time PCR. All plots indicate mean ± SD, and represent n=4 independent replicates.

**Figure S4. Related to Figure 4.**
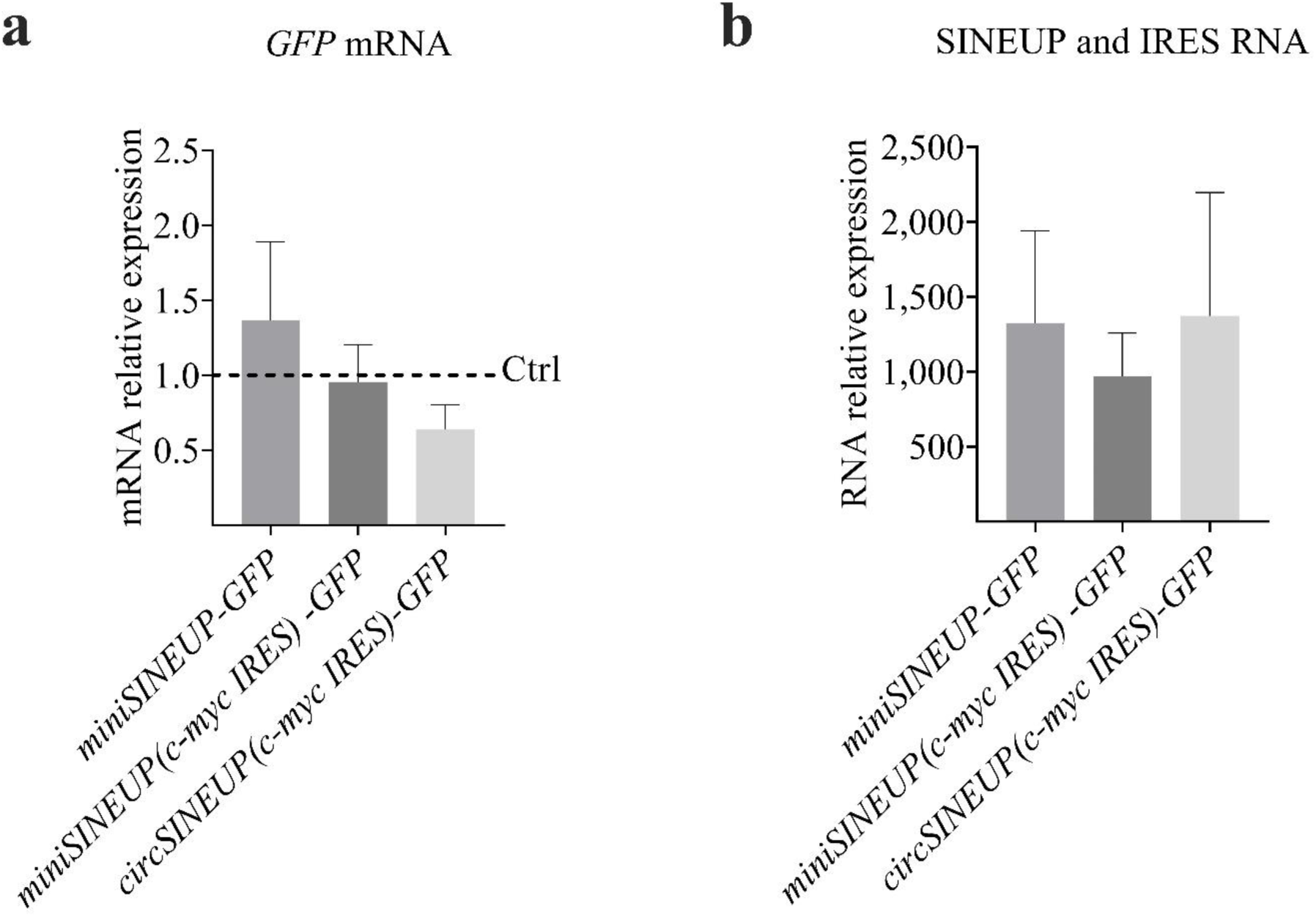
**Circularized SINEUP RNA maintains its activity in *trans*.** *circSINEUP(c-myc IRES)-GFP* activity. Experiments were performed as described in methods (“Cell Lines and Transfection Conditions”). HEK293T cells were co-transfected with circRNA-expressing vector and pEGFP (to express GFP protein) in a 6:1 ratio. 48 h after transfection, cells were collected and RNA extracted. Empty pCDNA3.1(-) was used as negative control (Ctrl=1, dotted line) and linear *miniSINEUP-GFP* and *miniSINEUP(c-myc IRES)-GFP* were the positive controls. RNAs were extracted, reverse transcribed and analyzed with real time PCR. RNA expressions. *GAPDH* was used to normalize. **a)** *GFP* mRNA levels. Empty pCDNA3.1(-) was the negative control (Ctrl=1, dotted line). *GFP* mRNA relative expressions were quantified and reported on the graph. No significant variations were observed. **b)** SINEUP and circSINEUP RNA. Expression of all transfected constructs were confirmed with real time PCR. All graphs indicate mean ± SD, and represent n=3 independent replicates.

**Figure S5. Related to Figure 5.**
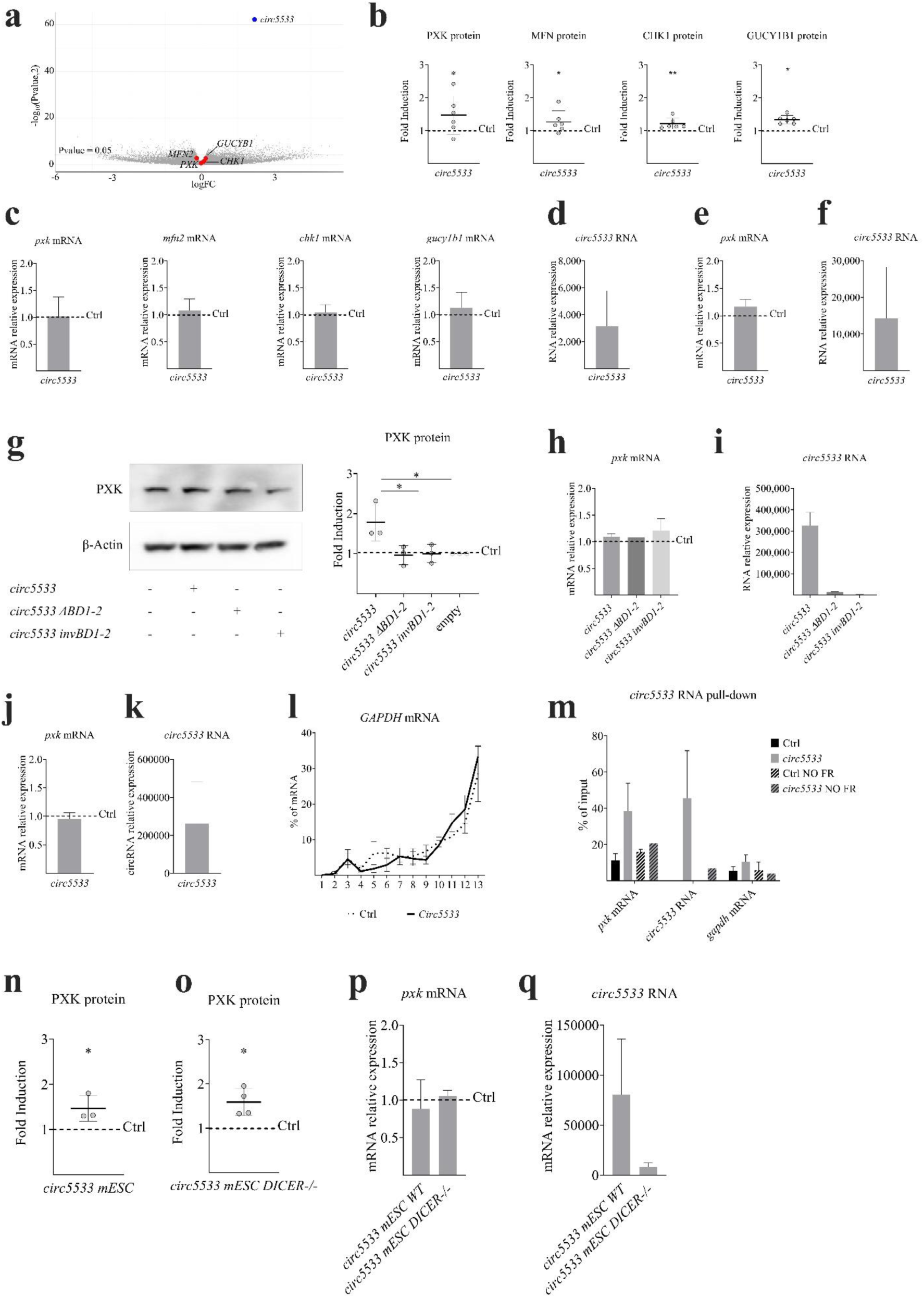
**(a-f) *circ5533* increases target protein levels without altering their mRNA levels, validation results.** HEK293T were transfected with pcDNA3.1(+) ZKSCAN1 MCS Exon *circ5533* (n=6). Control cells were transfected with empty pcDNA3.1(+) ZKSCAN1 MCS Exon Vector (as described in methods “Cell Lines and Transfection Conditions”). 48 h after transfection, cells were harvested; cell pellets were divided into two tubes in order to extract both proteins and RNA from the same sample. **a)** RNA-seq volcano plot. log_2_FC is reported on nthe x-axis while −log_10_ of the FDR-corrected p-values are plotted on the y-axis. Dots associated to negative values of log_2_FC represent downregulated genes, dots on the right represent upregulated genes. Red dots represent genes codifying for proteins whose fold inductions (FC>1.5) were validated with western blot. Expression levels of the genes of interest were validated with real time PCR (Figure S6c). Blue dot represents over-expressed *circ5533.* p <0.05 was considered significant. *Circ5533* levels were validated with real time PCR (Figure S6c). **b)** Mass spectrometry results validation. Total protein extract was analyzed with western blot. β-actin and GAPDH were used to normalize; fold-inductions were calculated relative to the empty pcDNA3.1(+) ZKSCAN1 MCS Exon Vector. PXK, MFN, CHK1, GUCY1B1 Abs were used to detect levels of proteins of interest. *circ5533* overexpression led to upregulation of all proteins of interest. Plots report protein fold induction mean ± SD; they represent 6 independent biological replicates; *p <0.05 and **p <0.01 were considered significant (one-sample t-test). **c)** *circ5533* target mRNA levels. Extracted RNAs were reverse-transcribed and analyzed with real time PCR. *GAPDH* was used to normalize. Empty pcDNA3.1(+) ZKSCAN1 MCS Exon Vector was the negative control (Ctrl=1, dotted line). Target mRNA relative expressions were quantified and reported on the graph. No significant variations were observed. **d)** *circ5533* RNA expression (from pcDNA3.1(+)ZKSCAN1 MCS Exon Vector). Overexpression of *circ5533* was observed with qRT-PCR using specific divergent primers, normalized on *GAPDH* mRNA levels and compared to the empty control. The graph indicates mean ± standard deviation (n=6). **e)** *Pxk* mRNA expression. As discussed in Figure 6c, HEK293T cells were transfected with pcDNA3.1(+)-Laccase2-circ5533 (n=9). Empty pcDNA3.1(+)-Laccase2 vector was the negative control. 48 h after transfection, cells were harvested and RNA extracted. Total RNA was reverse transcribed and analyzed with real time PCR. *GAPDH* was used to normalize. Empty pcDNA3.1(+)-Laccase 2 MCS Exon Vector was the negative control (Ctrl=1, dotted line). *Pxk* mRNA relative expressions were quantified and reported on the graph. No significant variations were observed. **f)** *circ5533*, from pcDNA3.1(+)-Laccase 2 vector, RNA expression. Over-expression of *circ5533* was observed with qRT-PCR using specific divergent primers, normalized on *GAPDH* mRNA levels and compared to empty pcDNA3.1(+)-Laccase 2 MCS Exon Vector. The graph indicates mean ± standard deviation (n=6). **(g-i) *Circ5533* mutants.** *circ5533* mutants were generated deleting both BD1 and 2 (*circ5533* ΔBD1-2) or inverting their sequence orientations (*circ5533* invBD1-2). HEK293T cells were transfected with *circ5533* mutants (n=3). Empty pcDNA3.1(+)-Laccase2 vector was the negative control while pcDNA3.1(+)-Laccase2-*circ5533* was the positive control. 48 h after transfection, cells were harvested and proteins and RNAs were extracted. **g)** Western blot analysis was performed, anti-PXK Ab was used to detect PXK protein. PXK fold induction was calculated compared to the empty control (Ctrl=1, dotted line). β-actin was used to normalize. Neither *circ5533* ΔBD1-2 nor *circ5533* invBD1-2 induced PXK expression. *circ5533* mutant activities were plotted as mean fold-induction values ± SD. *p <0.05 was considered significant. Ordinary one-way ANOVA, with Holm-Sidak’s multiple comparisons test, was used to calculate significance. **h)** Pxk mRNA expression. Total RNA was reverse-transcribed and analyzed with real time PCR. *GAPDH* was used to normalize. Empty pcDNA3.1(+)-Laccase 2 MCS Exon Vector was the negative control (Ctrl=1, dotted line). *Pxk* mRNA relative expressions were quantified and reported on the graph. No significant variations were observed. **i)** circRNA expression. Overexpression of circRNAs were observed with qRT-PCR using specific divergent primers, normalized on *GAPDH* mRNA levels and compared to empty pcDNA3.1(+)-Laccase 2 MCS Exon Vector. Both *circ5533* mutants were expressed at lower levels than WT *circ5533*. The graph indicates mean ± standard deviation (n=3). **(j-m) *circ5533*: a circSINEUP able to interact with *Pxk* mRNA inducing its protein translation.** HEK293T cells were transfected with pcDNA3.1(+)-Laccase2-*circ5533* (n=4). Empty pcDNA3.1(+)-Laccase2 vector was the negative control. (**j** and **k**) Following CHX treatment, 1/10 of the sample was used to perform total RNA extraction to assess both *Pxk* mRNA and *circ5533* expression levels. *GAPDH* was used to normalize. Empty pcDNA3.1(+)- Laccase was the negative control (Ctrl=1, dotted line). **l)** As described in Figures 1d and 6d, lysates from control and treated cells were fractionated by sucrose gradients and separated through an UV detector reading the RNA amount (254 nm) in each fraction. The relative distributions (in %) of *GAPDH* mRNA were analyzed by RT-qPCR in each of the 13 gradient fractions for both control and treated samples. RNA distribution was reported as the mean ± SD of 3 replicates. *Circ5533* RNA did not affect *GAPDH* mRNA distribution. **m)** *circ5533* RNA Pull Down (RPD). *Circ5533* was purified using biotinylated DNA probes whose length ranges around 20 nts. As negative control, RPD was performed in the absence of targeting probes. qRT-PCRs show the enrichment of *circ5533* and *Pxk* mRNA upon *circ5533* RPD. Data are shown as means of both *circ5533* and *Pxk* mRNA enrichment versus input relative to the negative control ± SD of 6 replicates. **(n and o) *circ5533* increases PXK protein levels in both mESC WT and mESC *Dicer^-/-^* cells.** *Pxk-flag* cDNA was co-transfected with *circ5533*(1:12) in wild type mouse embryonic stem cells (mESC) or *Dicer* knock-out mESCs. After 48 h, cells were harvested and proteins and RNA were extracted. Empty pcDNA3.1(+)-Laccase2 vector, cotransfected with *Pxk*-*flag*, was used as negative controls; *Pxk*-*flag* WT + *circ5533* was the positive control. **(n)** Western blot analysis was performed, anti-PXK Ab was used to detect PXK protein. PXK fold induction was calculated compared to the empty control (Ctrl=1, dotted line). β-actin was used to normalize. *circ5533* induced PXK expression in both WT and *Dicer^-/-^* mESC . *circ5533* activity was plotted as mean fold-induction values ± SD. *p <0.05 was considered significant. Ordinary one-way ANOVA, with Holm-Sidak’s multiple comparisons test, was used to calculate significance. **(p)** *Pxk* mRNA relative expression. *GAPDH* was used to normalize. Empty pcDNA3.1(+)-Laccase 2 cotransfected with *Pxk-flag* was the negative control (Ctrl=1, dotted line). *Pxk* mRNA relative expressions were quantified and reported on the graph. No significant variations were observed. **(q)** Over-expression of circRNAs was observed with qRT-PCR using specific divergent primers, normalized on *GAPDH* mRNA levels and compared to empty pcDNA3.1(+)-Laccase 2 MCS Exon Vector. Graph indicates mean ± standard deviation (n=3).

**Figure S6. Related to Figure 6.**
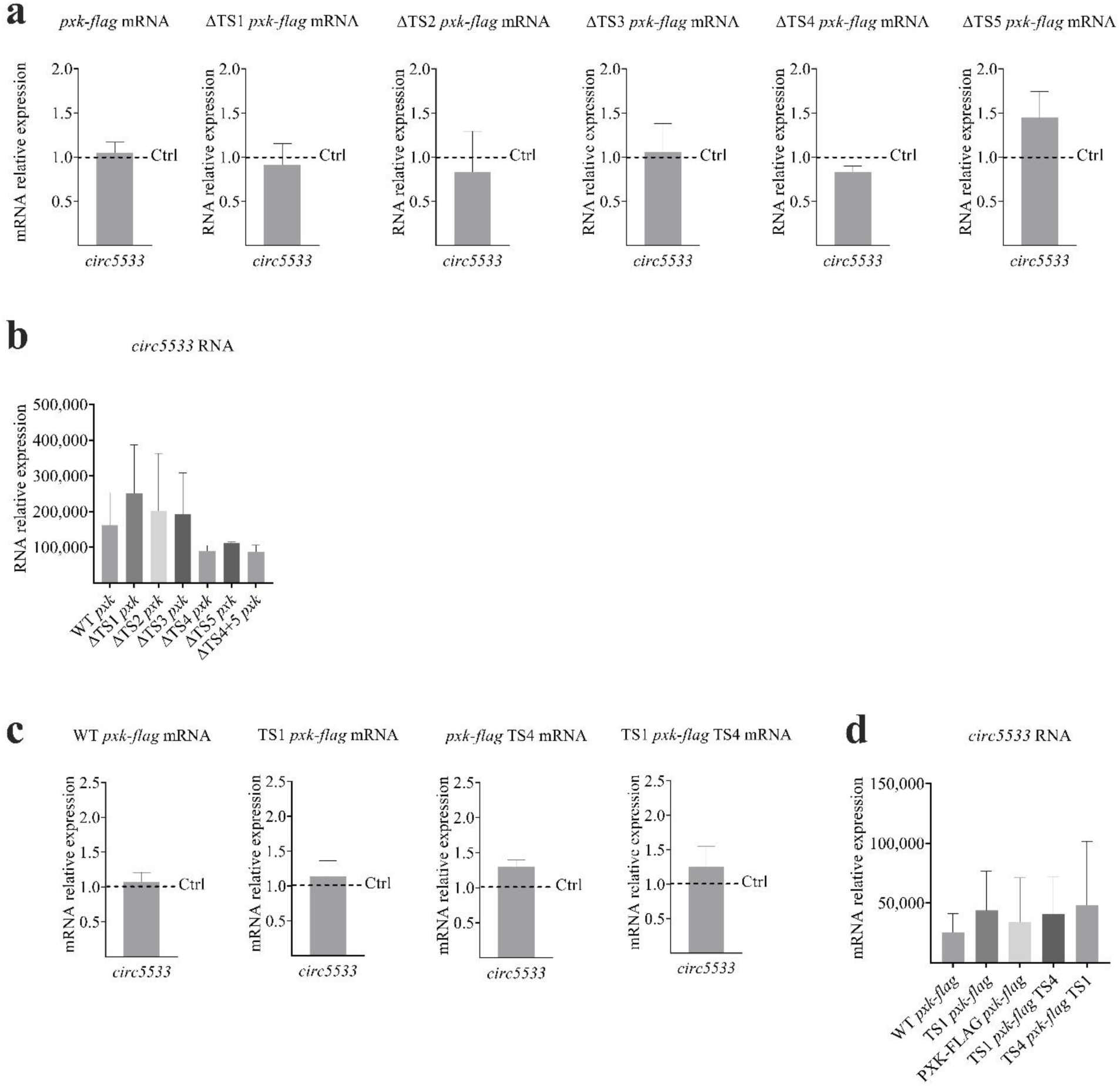
***Pxk-flag* mutants RNA.** Each *Pxk* mutant construct was cotransfected with *circ5533* in HEK293T cells (1:12). After 48 h, cells were harvested and proteins and RNA were extracted. Empty pcDNA3.1(+)-Laccase2 vector cotransfected with different *Pxk*-mutants were used as negative controls; *Pxk-flag* WT + *circ5533* was the positive control. **a)** ΔTSs *Pxk* mutant mRNA. RNA was extracted, reverse-transcribed and analyzed with qRT-PCR. *GAPDH* was used to normalize. Empty pcDNA3.1(+)-Laccase 2 cotransfected with each *Pxk* mutant was the negative control (Ctrl=1, dotted line). *Pxk* mRNA relative expressions were quantified and reported on the graph. No significant variations were observed. **(b and d)** circRNA expression. Overexpression of circRNAs was observed with qRT-PCR using specific divergent primers, normalized on *GAPDH* mRNA levels and compared to empty pcDNA3.1(+)-Laccase 2 MCS Exon Vector. Graph indicates mean ± standard deviation (n=3). **c)** TSs *Pxk* mutant mRNA. Total RNA was extracted at 48 h after cotransfection, reverse-transcribed and analyzed with qRT-PCR. *GAPDH* was used to normalize. Empty pcDNA3.1(+)-Laccase 2 co-transfected with each *Pxk* mutant was the negative control (Ctrl=1, dotted line). *Pxk* mRNA relative expressions were quantified and reported on the graph. No significant variations were observed.

**Table S1.** List of IRESes tested in SINEUP-RNAs.

| Origin | IRES type | Size (bp) | Ref number<br>(IRESite Id) |
| --- | --- | --- | --- |
| Viral | HCV | 383 | 222 |
|  | Poliovirus | 312 | 242 |
|  | EMCV | 576 | 140 |
|  | CrPV | 192 | 40 |
| Cellular | c-myc | 41 & 395 | 35 |
|  | Apaf1 | 231 | 110 |
|  | ELG1 | 460 | 492 |
|  | DMD | 71 |  |

**Table S2.**
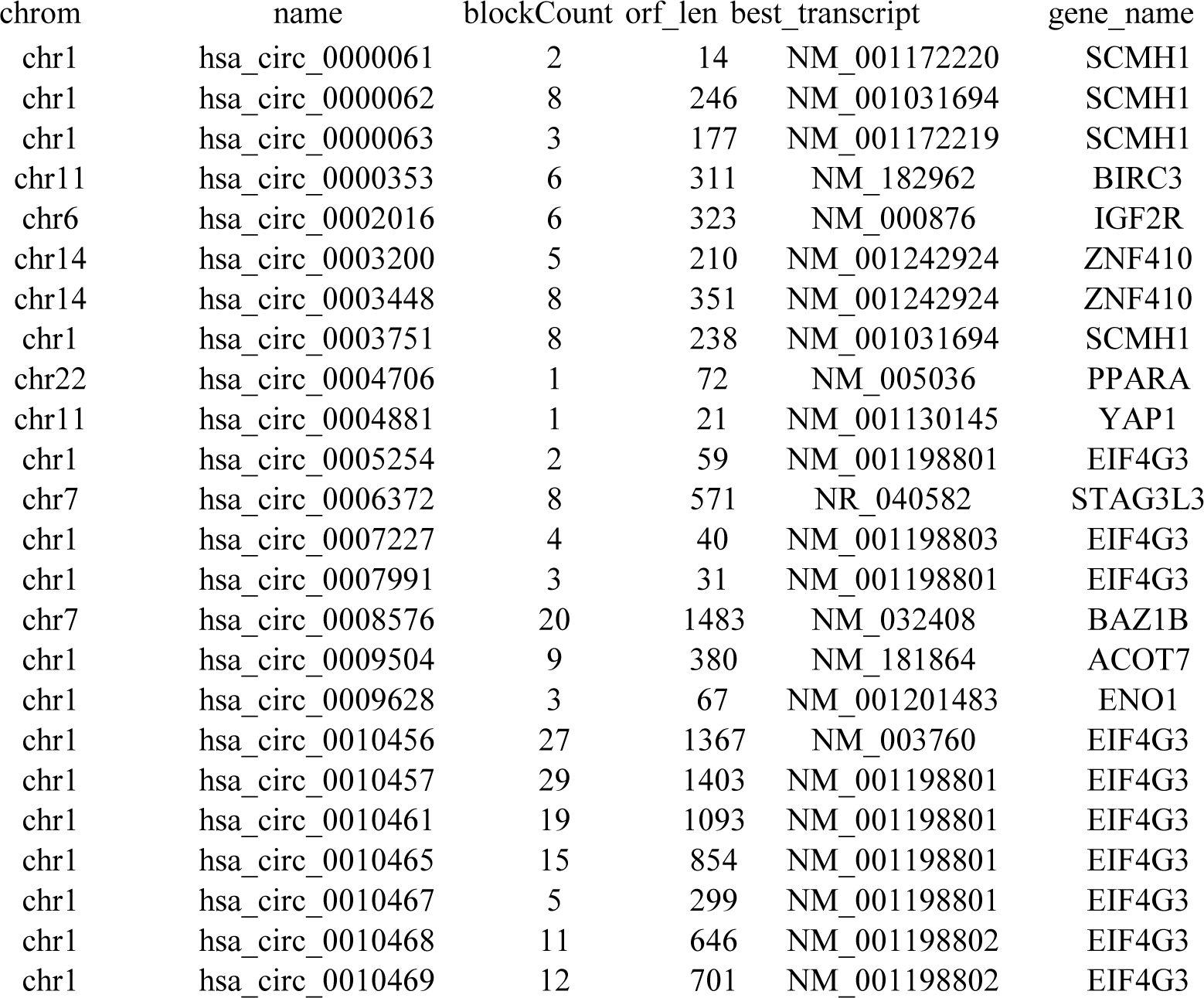
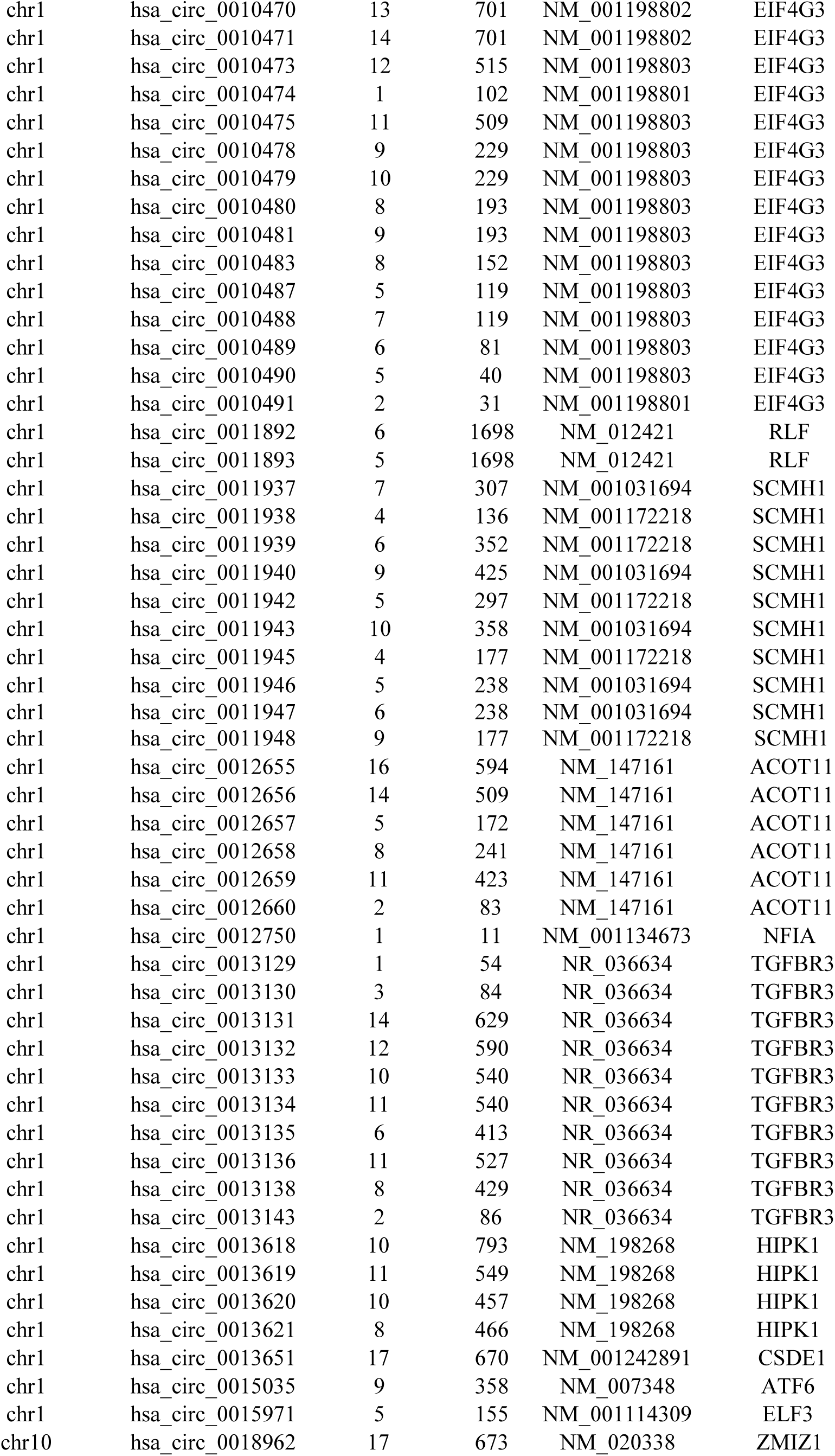
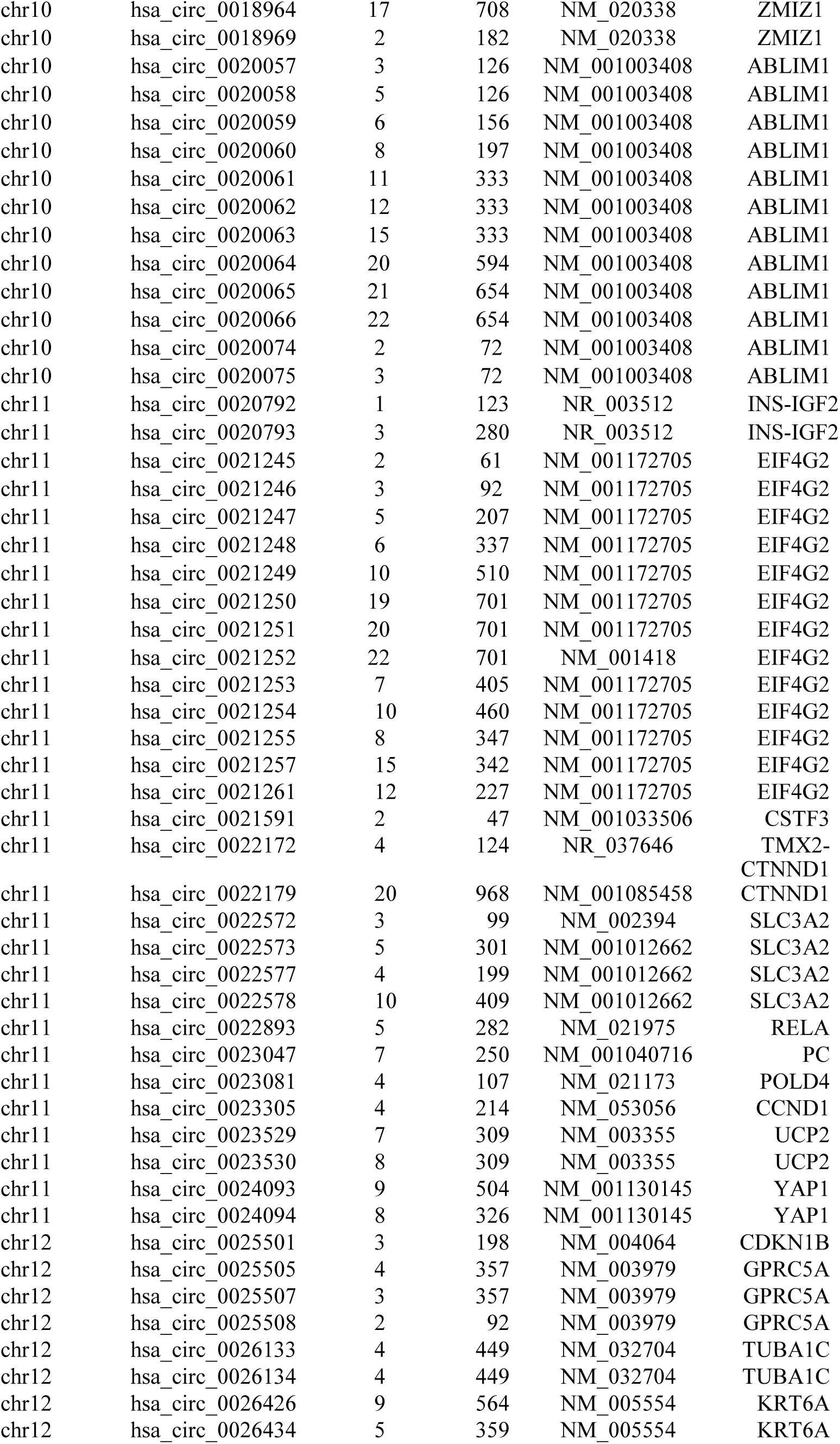
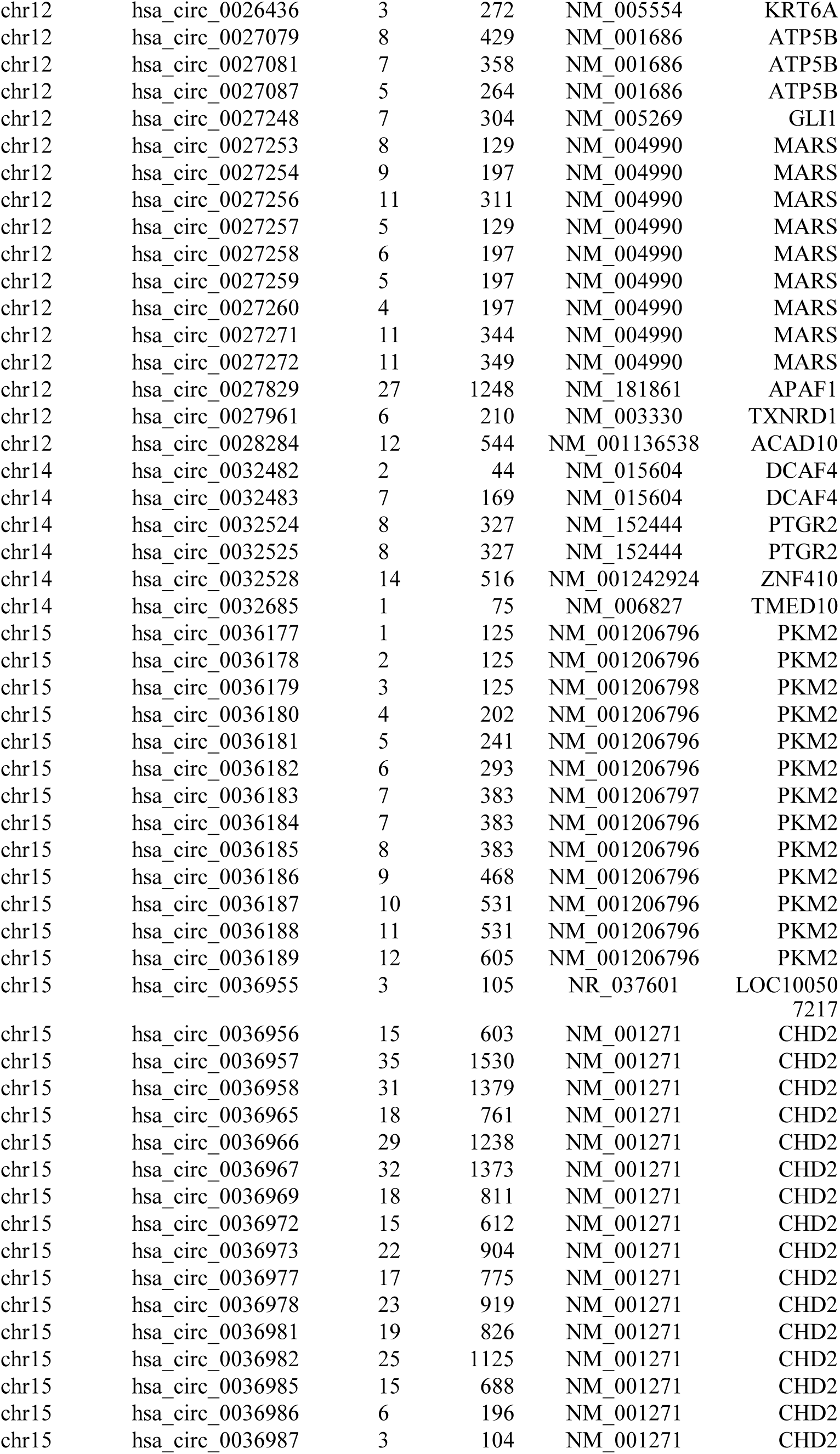
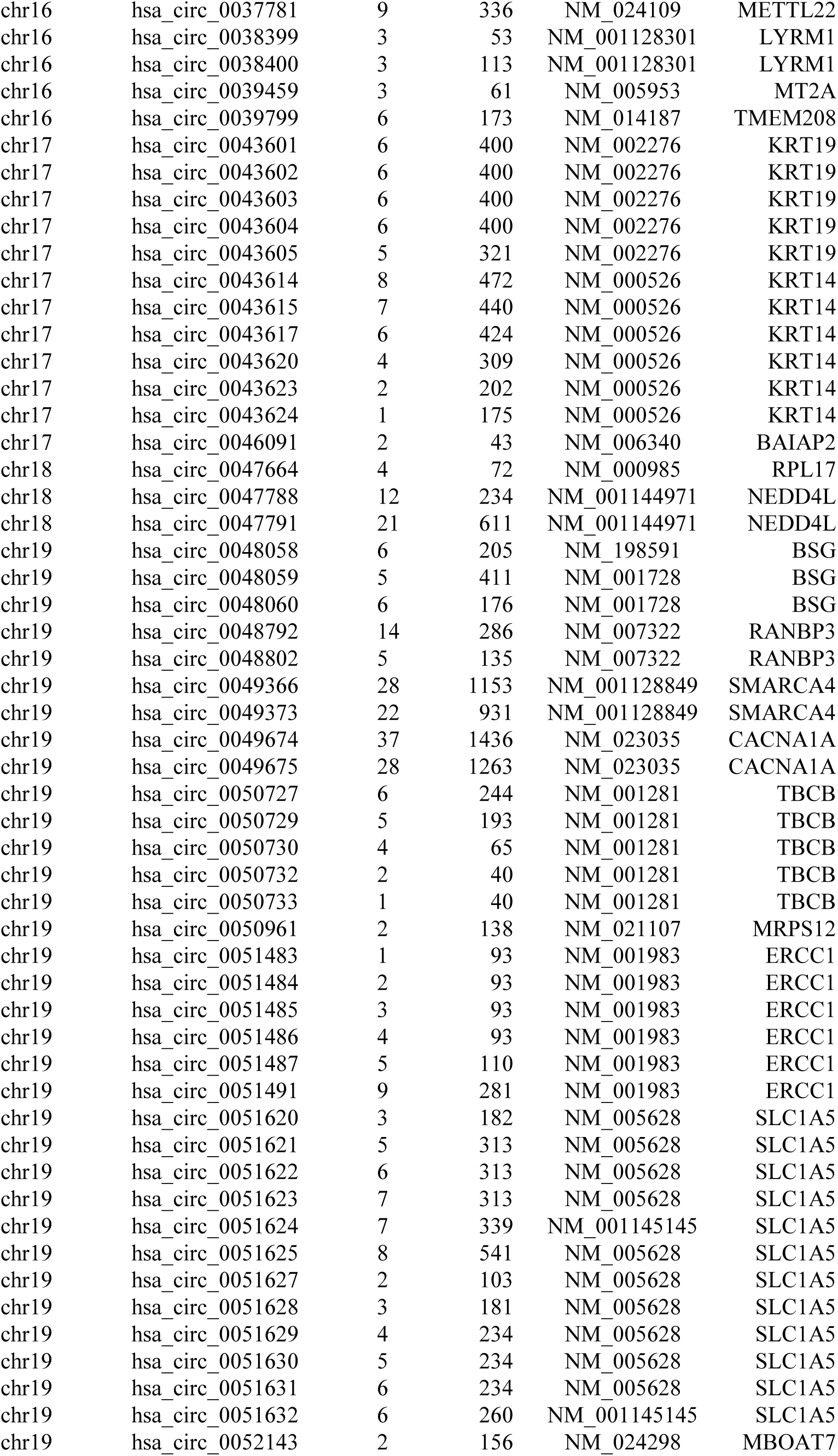
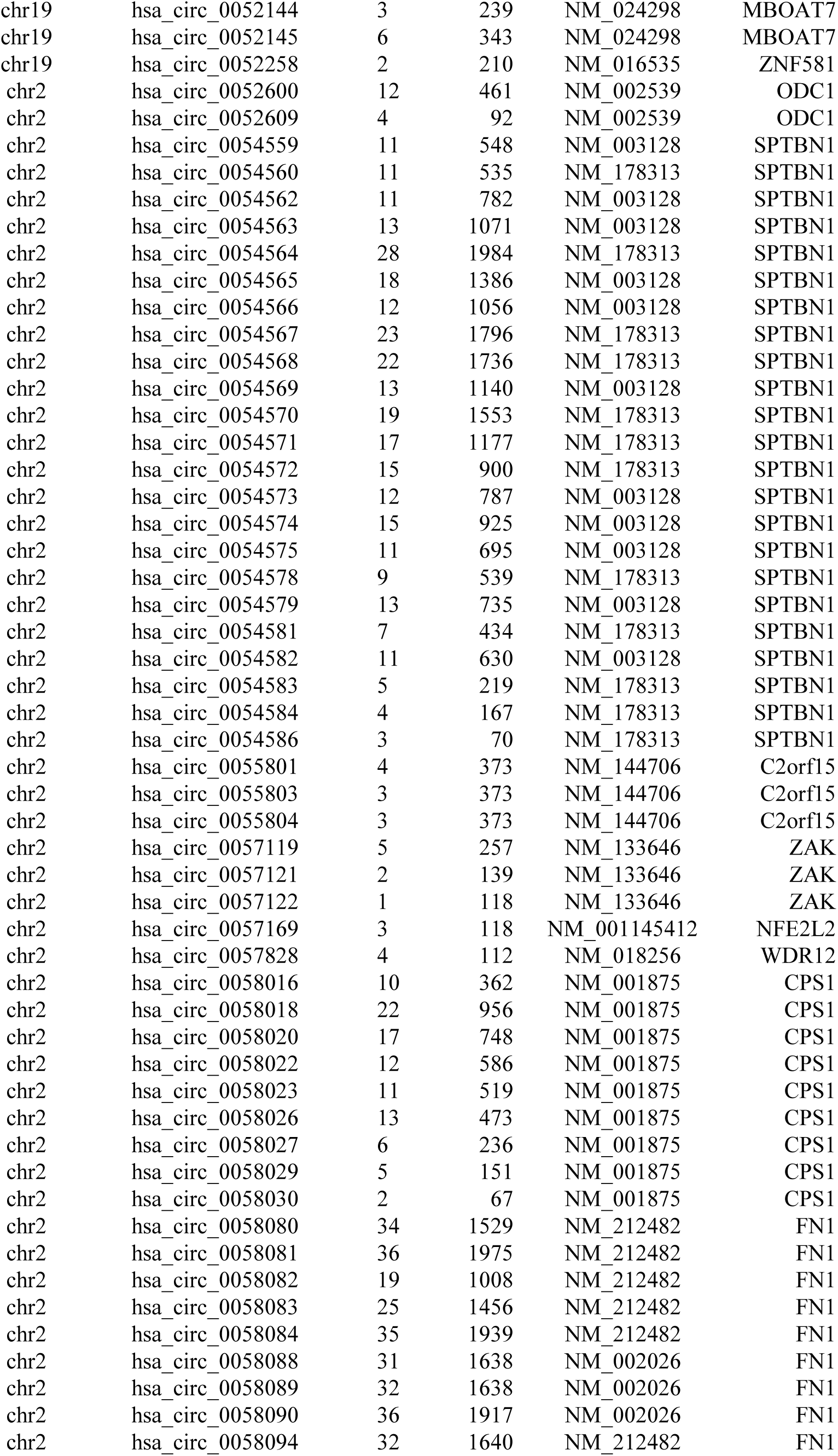
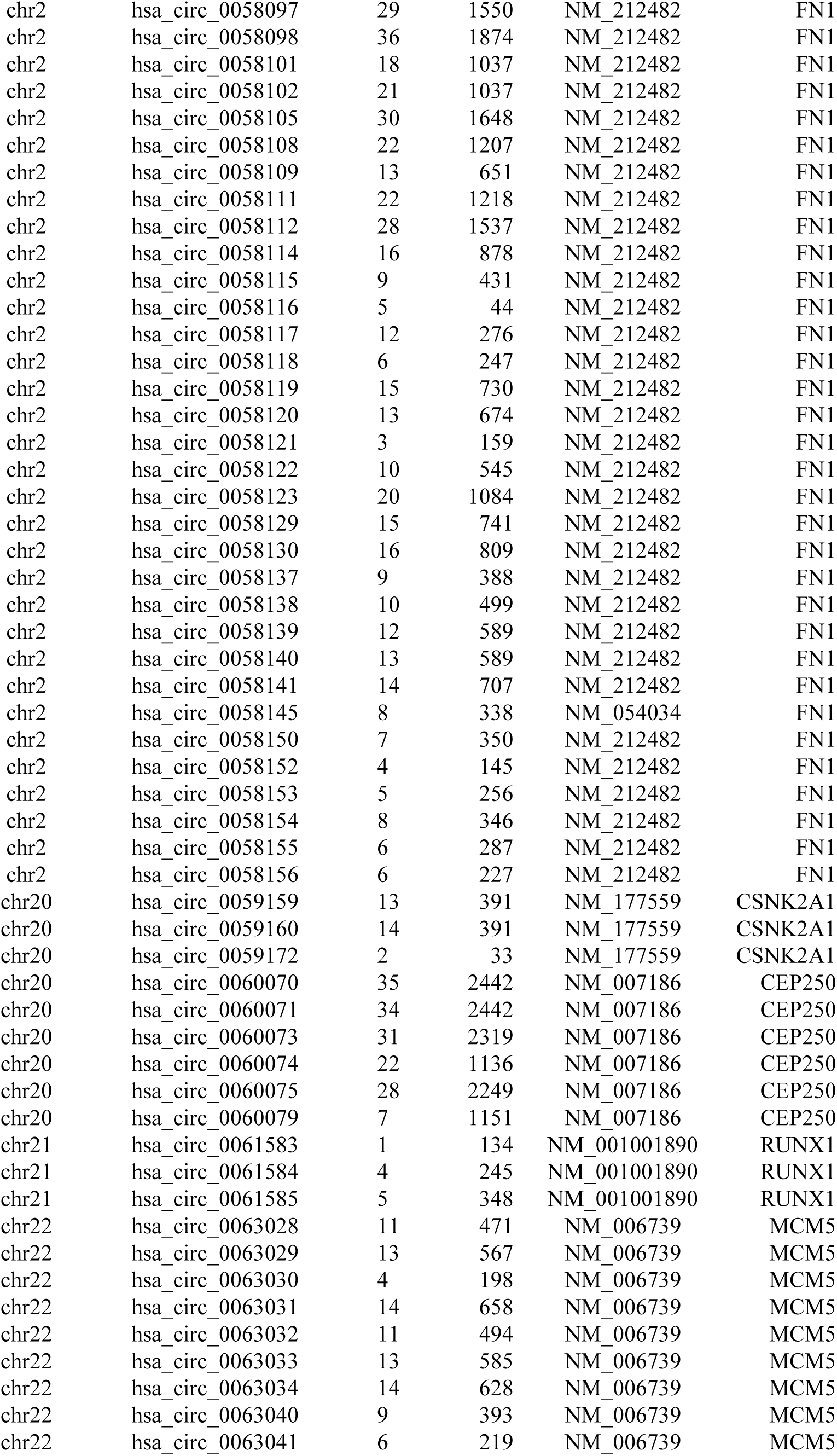
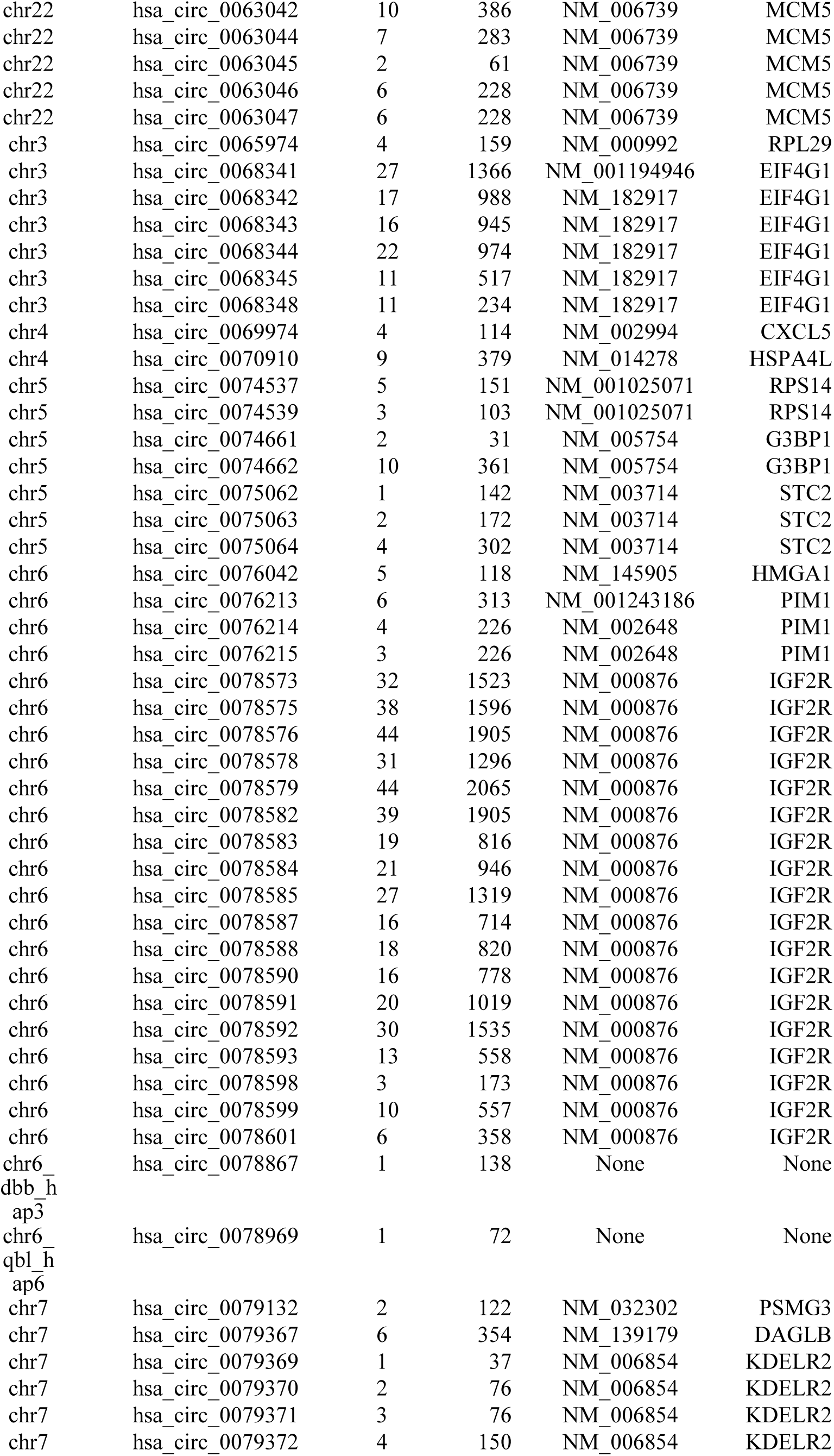
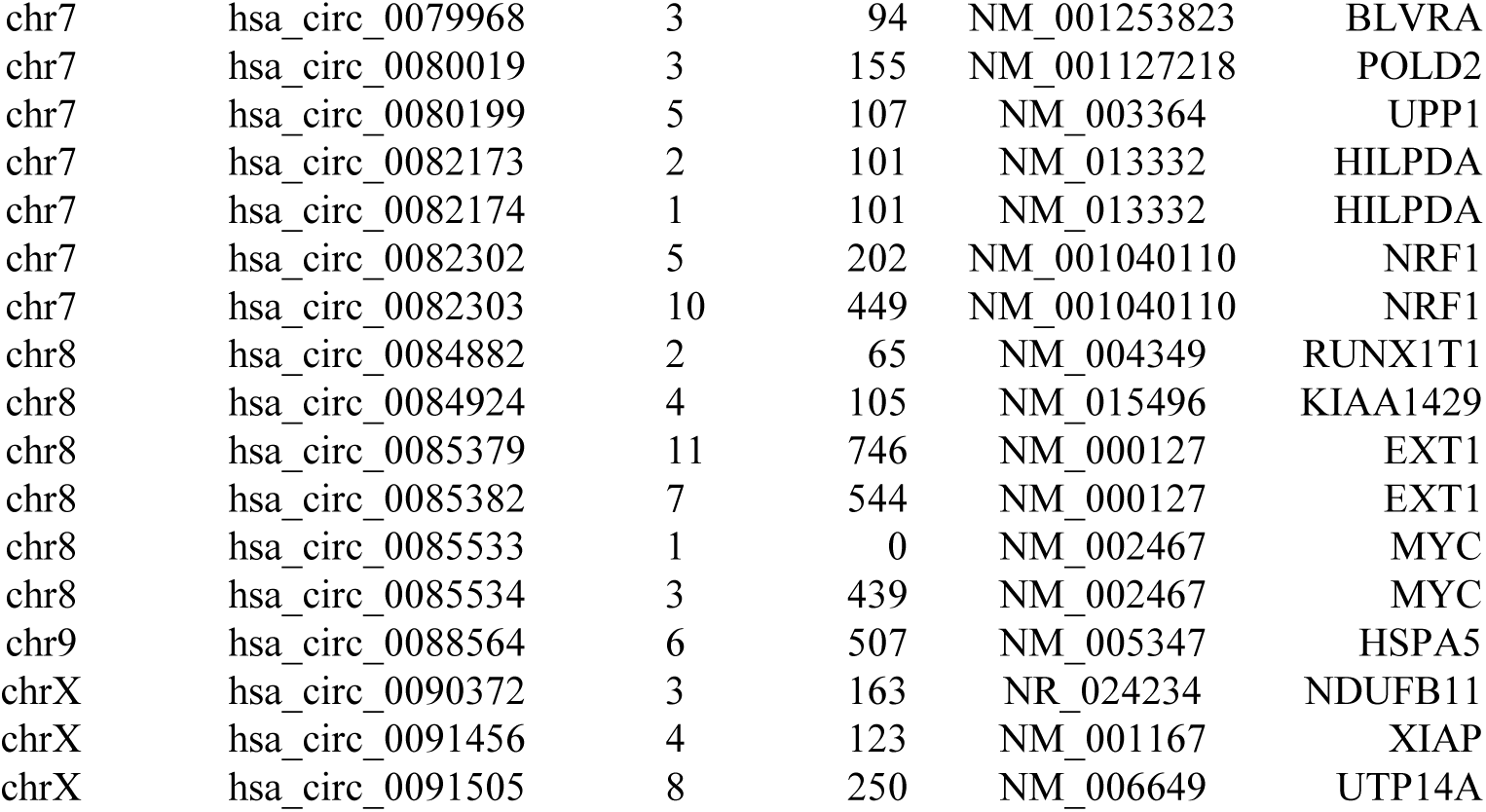
Circular RNAs with potential IRES activity.

**Table S3.**
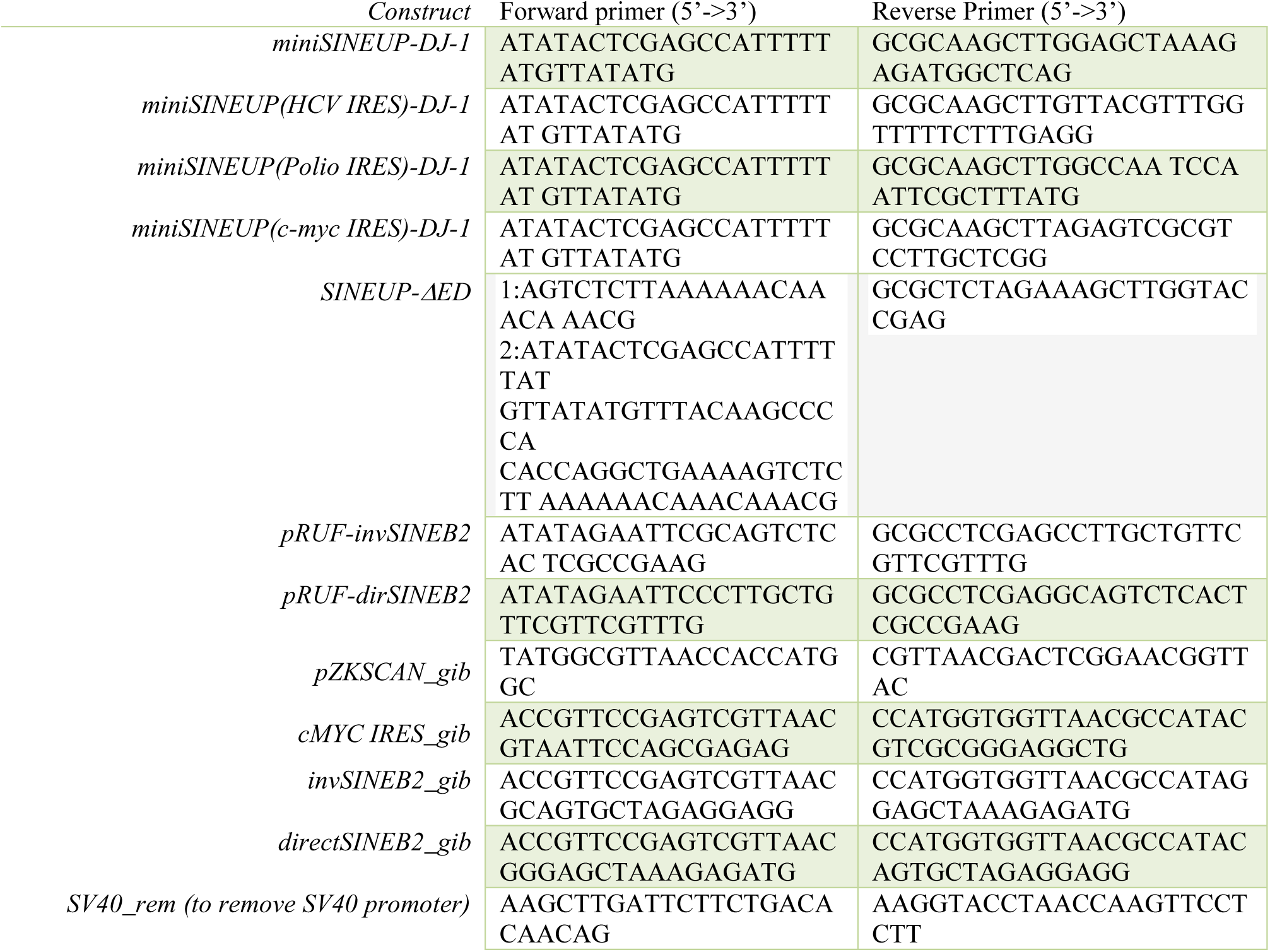
Sequence of primers used in the miniSINEUP-DJ, miniSINEUP-IRES-DJ1, miniSINEUP-(HCV IRES)-GFP and pRUF-SINEB2 (direct and inverted) clones.

**Table S4.**
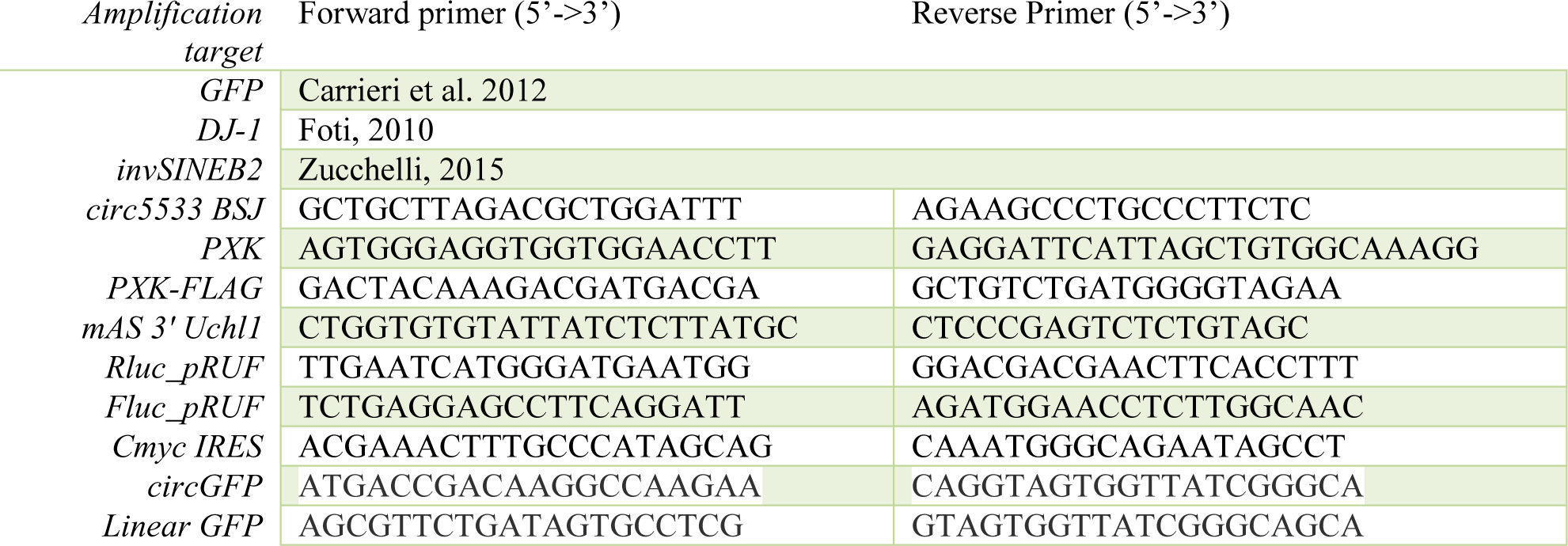
List of primers used for qRT-PCR Real-time analysis.

**Table S5.** List of antibodies.

| Antibody | Source | Identifier |
| --- | --- | --- |
| *anti-DJ-1* | Enzo Life Technology | ADI-KAM-SA100-E |
| *anti-β-Actin* | Sigma | A2066 |
| *anti-PXK* | Sigma | HPA024068-100UL |
| *anti-MFN2* | Cell Signaling | 9482S |
| *anti-CHK1* | Proteintech | 25887-1-AP |
| *anti-GUCY1B1* | Abcam | ab154841 |
| *anti-FLAG* | Sigma | F3165 |

